# Dramatic Changes in Mitochondrial Subcellular Location and Morphology Accompany Activation of the CO_2_ Concentrating Mechanism

**DOI:** 10.1101/2024.03.25.586705

**Authors:** Justin Findinier, Lydia-Marie Joubert, Michael F. Schmid, Andrey Malkovskiy, Wah Chiu, Adrien Burlacot, Arthur R. Grossman

## Abstract

Dynamic changes in intracellular ultrastructure can be critical for the ability of organisms to acclimate to environmental conditions. Microalgae, which are responsible for ∼50% of global photosynthesis, compartmentalize their Rubisco into a specialized structure known as the pyrenoid when the cells experience limiting CO_2_ conditions; this compartmentalization appears to be a component of the CO_2_ Concentrating Mechanism (CCM), which facilitates photosynthetic CO_2_ fixation as environmental levels of inorganic carbon (Ci) decline. Changes in the spatial distribution of mitochondria in green algae have also been observed under CO_2_ limiting conditions, although a role for this reorganization in CCM function remains unclear. We used the green microalgae *Chlamydomonas reinhardtii* to monitor changes in the position and ultrastructure of mitochondrial membranes as cells transition between high CO_2_ (HC) and Low/Very Low CO_2_ (LC/VLC). Upon transferring cells to VLC, the mitochondria move from a central to a peripheral location, become wedged between the plasma membrane and chloroplast envelope, and mitochondrial membranes orient in parallel tubular arrays that extend from the cell’s apex to its base. We show that these ultrastructural changes require protein and RNA synthesis, occur within 90 min of shifting cells to VLC conditions, correlate with CCM induction and are regulated by the CCM master regulator CIA5. The apico-basal orientation of the mitochondrial membrane, but not the movement of the mitochondrion to the cell periphery, is dependent on microtubules and the MIRO1 protein, which is involved in membrane-microtubule interactions. Furthermore, blocking mitochondrial electron transport in VLC acclimated cells reduces the cell’s affinity for inorganic carbon. Overall, our results suggest that CIA5-dependent mitochondrial repositioning/reorientation functions in integrating cellular architecture and energetics with CCM activities and invite further exploration of how intracellular architecture can impact fitness under dynamic environmental conditions.

## INTRODUCTION

Both the intracellular position and ultrastructure of organelles can exhibit dynamic behavior in response to specific stressors or during metabolic shifts (1). Changes in spatial and morphological features of an organelle may be key for completing developmental processes and acclimating to changing environmental conditions. For example, the movement and repositioning of mitochondria in the cell can be required for their proper partitioning between mother and daughter cells in budding yeast (2). They are also critical in animals for optimizing delivery of energy for fueling cell migration (3, 4) and the release of synaptic neurotransmitters (5), and generally require the activity of the cytoskeletal network (6). The movement and repositioning of cellular compartments have also been demonstrated for the ER (7, 8), nuclei (9, 10) and chloroplasts (11).

In microalgae, which mediate ∼50% of photosynthesis on Earth (12), major cellular ultrastructural shifts can occur as CO_2_ levels change. One change reflects the condensation of most of the CO_2_-fixing enzyme Ribulose-1,5-bisphosphate carboxylase (Rubisco) in a membrane-less organelle called the pyrenoid (13). Such a structure is associated with the induction of a CO_2_ concentrating mechanism (CCM), which elevates the cell’s affinity for inorganic carbon (Ci, which includes CO_2_, HCO_3_^-^, CO_3_^-2^) by actively concentrating CO_2_ in the pyrenoid matrix (14).

Dramatic alterations in ultrastructure and intracellular positioning of mitochondria in response to changing CO_2_ levels have also been reported (15–17). In the model green microalga *Chlamydomonas reinhardtii* (Chlamydomonas hereafter), the mitochondrial membranes are located mostly within the ‘cup’ formed by the single chloroplast when the cells are grown under conditions of high CO_2_ availability (HC, 2-5% CO_2_ in air, or in the presence of acetate which drives high respiratory CO_2_ production). In contrast, as CO_2_ availability declines because of a limited supply or increased consumption, the mitochondrial membranes move to the cell periphery and become wedged between the chloroplast outer envelope and the plasma membrane (15, 17). While the functional consequences of these mitochondrial structural modifications remain elusive, their correlation with CCM induction suggests a role in Ci acquisition.

In Chlamydomonas, the major CCM components include carbonic anhydrases (CAHs), and HCO_3_^-^ and CO_2_ transporters. They enable the cells to efficiently transport Ci and increase the CO_2_ concentration in the pyrenoid (18) where most of the Rubisco resides. Despite evidence that CCM activation is a continuum tightly regulated by CO_2_ availability (19), distinct modes of Ci transport are generally considered to operate at: (i) HC, (ii) ambient or low CO_2_ (LC, 0.04% CO_2_), and (iii) very low CO_2_ (VLC, <0.02% CO_2_) (**Fig. 1**). In HC, the CCM is inactive, and CO_2_ passively diffuses across cellular membranes and into chloroplasts where it is fixed by Rubisco. In LC, most CO_2_ diffuses into the cell, is converted into HCO_3_^-^ and trapped in the chloroplast stroma through the activity of LCIB (20, 21). At VLC, the cells mainly actively transport HCO_3_^-^ across the plasma membrane by the HLA3 transporter (22) and move it into the chloroplast stroma through the inner envelope membrane channel LCIA (21, 23). In LC and VLC, stromal HCO_3_^-^ is transported to the thylakoid lumen by the bestrophin-like transporters, BST1-3 (24), and is then converted back to CO_2_ by CAH3 in the thylakoid tubules that penetrate the pyrenoid (25). The CO_2_ then diffuses into the pyrenoid matrix where it is fixed by Rubisco. Also at VLC, the CCM protein LCIB encases the pyrenoid, potentially capturing CO_2_ that leaks from the pyrenoid (converting it back to HCO_3_^-^) (26). Acclimation of Chlamydomonas to LC and VLC depends on CIA5, considered the CCM master regulator (27, 28), and CAS1, a pyrenoid localized calcium sensor protein (29). Mutations that eliminate either of these proteins block the induction of most genes associated with LC and VLC responses (29–32).

**Figure 1.**
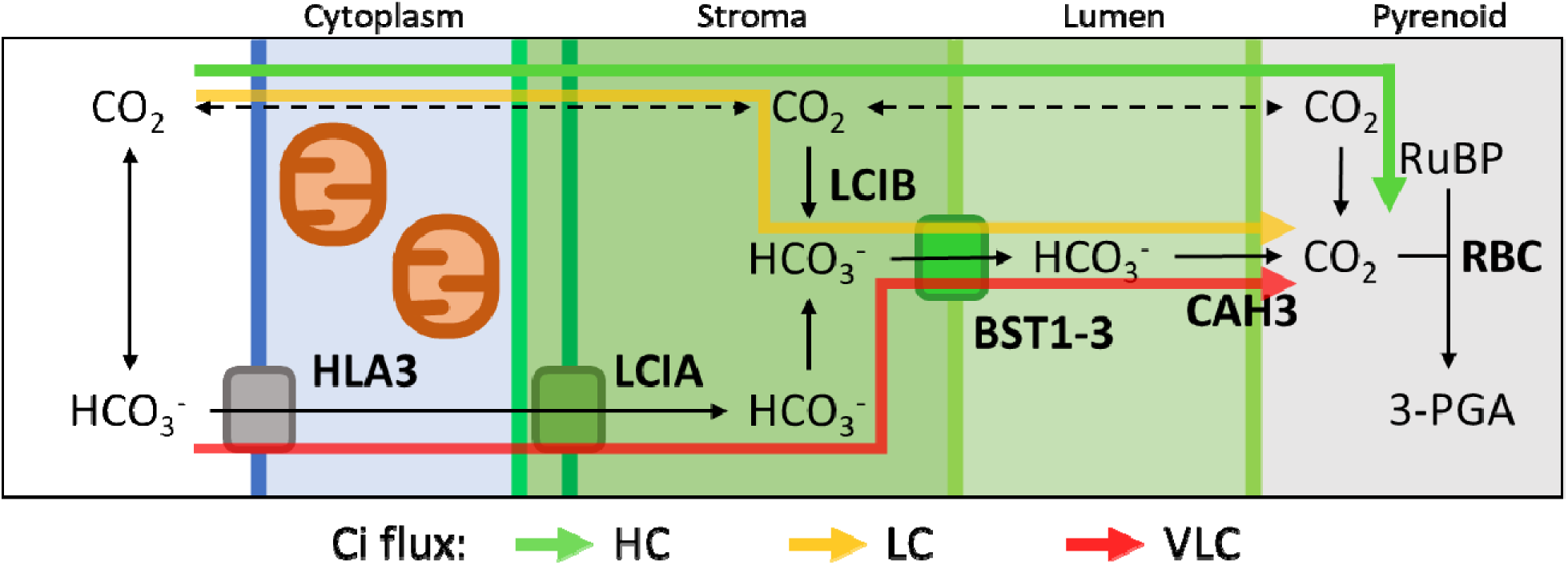
Modes of CO_2_ delivery to Rubisco. At HC, CO_2_ passively diffuses to Rubisco (green arrow). At LC, Ci is trapped in the stroma as HCO_3_^-^, which is facilitated by LCIB activity (yellow arrow). At VLC, HCO_3_^-^ is directly taken up from the medium by the plasma membrane transporter HLA3 and then channeled from the cytoplasm to the stroma by LCIA (red arrow). Stromal HCO_3_^-^ is transported into the thylakoid lumen by BST1-3 and delivered as CO_2_ to the pyrenoid through CAH3 activity. Acronyms: CO_2_: Carbon dioxide; HCO_3_^-^: bicarbonate; HLA3: High Light-Activated 3; LCIA: Low CO_2_ Induced A; LCIB: Low CO_2_ Induced B; BST1-3: Bestrophin 1-3; CAH3: Carbonic Anhydrase 3.

In vascular plants, metabolic interactions between mitochondria and chloroplasts play a central role in maintaining the proper functioning of chloroplasts under various conditions (33, 34). In Chlamydomonas there are hints that mitochondria assume a new intracellular position and support CCM function when the cells experience low levels of Ci. RNAi studies showed that CCM activity is impacted by low levels/absence of the mitochondria-localized CAH4/5 proteins and CCP1/2 transporters (35, 36). The authors proposed that the mitochondrial CAHs and potential HCO_3_^-^ transporters encoded by CCP1/2 could recapture CO_2_ generated by mitochondrial respiration, photorespiration, and/or leakage from the chloroplast by converting it to HCO_3_^-^, allowing it to be shuttled back into the chloroplast. Mitochondrial respiration has also been proposed to help energize plasma membrane Ci uptake, likely by using reductant generated by chloroplast-to-mitochondria electron flow (37, 38). While mitochondrial relocation has been proposed to play a role in CCM function, it remains unclear to which CCM mode of action it is linked, the molecular mechanisms associated with this dramatic rearrangement, and the specific functions it provides for concentrating Ci.

In this study, we used a mitochondria-targeted fluorophore to monitor mitochondrial relocation as Chlamydomonas cells transition between HC, LC, and VLC conditions. Within 90 min of a transition from HC to VLC, the mitochondria relocate to the cell’s cortex with alignment of tubular mitochondrial membranes along the apico-basal axis. This alignment is strongly disrupted when microtubule formation is inhibited and by disruption of a gene encoding a homolog of a microtubule/mitochondrion interacting protein (MIRO1). This dynamic relocation is shown to be under the control of the CCM regulator CIA5 and correlates with relocation of LCIB from a diffuse stromal distribution to being concentrated around the pyrenoid, the induction of CCM genes, including *HLA3* and *CAH4*, and an increase in the cell’s affinity for Ci. Furthermore, we show that this change in Ci affinity is inhibited when VLC-maintained cells are exposed to respiratory inhibitors or have a genetic background in which the *CAH4* and *CAH5* genes were disrupted, but not when the apico-basal orientation of the tubular mitochondrial membranes was affected (e.g. in presence of inhibitors of microtubule formation or absence of MIRO1). Overall, our results highlight the kinetic and morphological features of mitochondrial relocation as cells transition to VLC conditions and suggest that the VLC mode of CCM function requires both energy generation by mitochondria and the interconversion of CO_2_ and HCO_3_^-^ within the mitochondria.

## RESULTS

### Dynamics of the mitochondrial network depends on CO_2_ availability

To investigate dynamic changes of the mitochondrial network, we generated a strain expressing the GFP variant fluorophore Clover targeted to the mitochondrial matrix (39). The spatial distribution of the network of mitochondrial tubules was then visualized by confocal microscopy (**Fig. 2**). When cells were grown photoautotrophically (TP medium) in moderate light, sparging cultures with HC or LC resulted in mitochondria mostly positioned within the cup formed by the chloroplast (**Fig. 2A**). Sparging cells with VLC caused the mitochondria to relocate to the cell periphery, between the chloroplast outer envelope and plasma membranes, where they appear as isolated dots and small tubules (**Fig. 2A**). To confirm the physiological state of the cells under the different conditions, we monitored, in a separately generated strain, the position of a fusion protein of the fluorophore mCherry with LCIB, a known marker that reports acclimation of cells to HC/LC and VLC (26). LCIB is diffuse throughout the chloroplast stroma in HC/LC, and only localizes to the perimeter of the pyrenoid in VLC, the same condition that results in mitochondrial localization to the cell periphery (**Fig. 2B**). These results indicate that mitochondrial relocation occurs in VLC (not HC or LC) acclimated state. We also monitored mitochondrial positioning under mixotrophic conditions (TAP medium, **Fig. 2C**) where acetate metabolism drives a higher respiration rate and an elevated level of internal CO_2_. The peripheral localization of the mitochondria was readily observed in TAP-grown cells when they were exposed to HL (conditions that drive CO_2_ consumption) or sparged with VLC, but not in cells exposed to low light (LL) (**Fig. 2C**). Furthermore, under conditions in which the mitochondria were peripherally located, most of the LCIB moved to the perimeter of the pyrenoid (**Fig. 2D**). These results further confirmed the specificity of the movement of the mitochondria to the cell periphery under VLC conditions.

**Figure 2.**
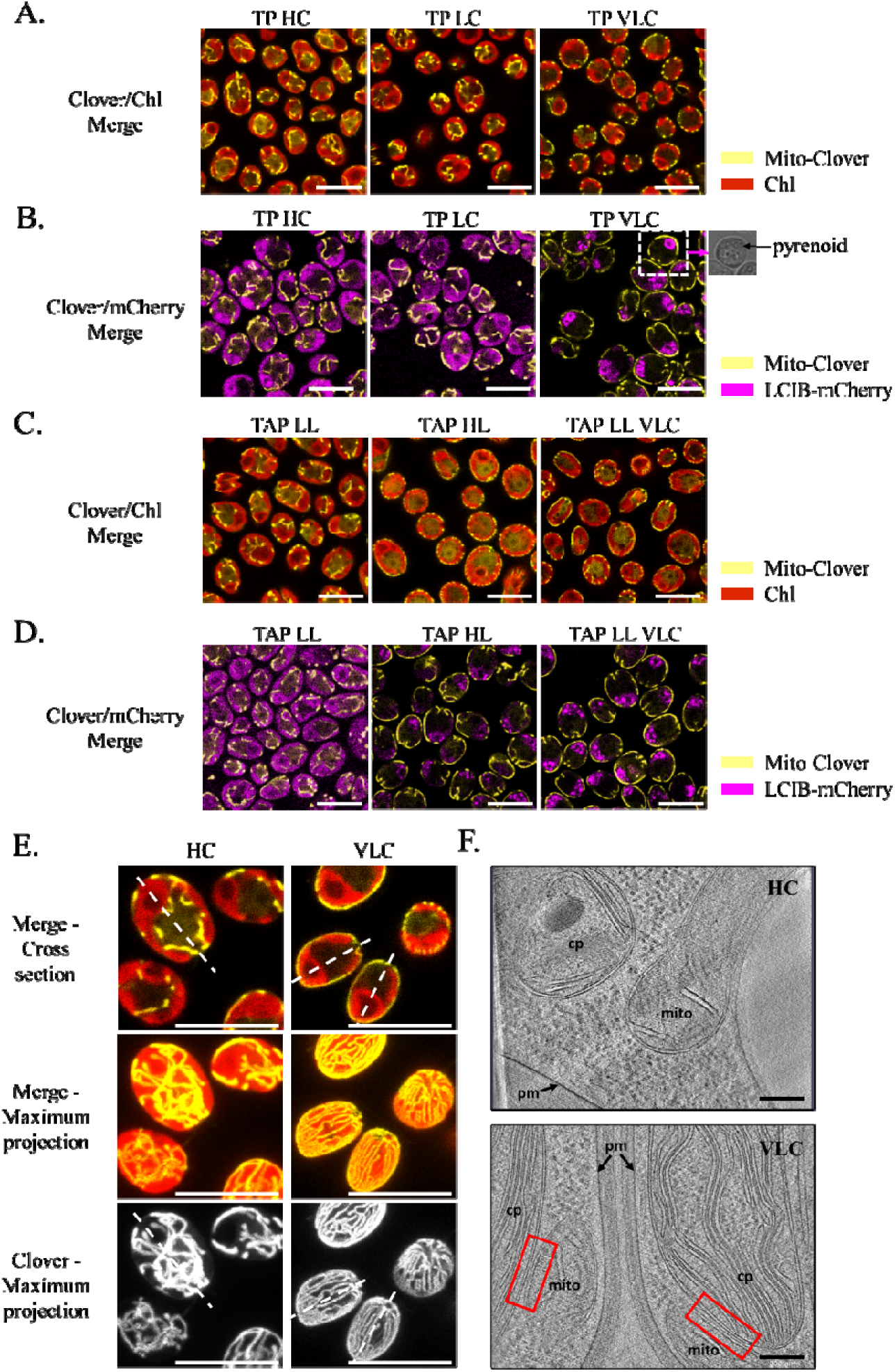
Physiological conditions and mitochondrial ultrastructure. **A.** Photoautotrophically grown cells were sparged with HC (∼2% in air), LC (0.04%), or VLC (< 0.02%) in Tris-Phosphate (TP) medium. Chlorophyll autofluorescence (red) marks the chloroplast while the Clover signal (yellow) marks the positions of the mitochondrial membranes. Scale bar: 10 µm. **B.** Localization of a LCIB-mCherry fusion (magenta) is monitored together with the mitochondria (yellow) in cells sparged with HC, LC, or VLC. Fluorescence from chlorophyll (Chl) was filtered out. The pyrenoid is shown in brightfield for one cell. Scale bar: 10 µm. **C.** Mitochondria localization was monitored in cells grown without aeration (shaken in flask, 120 rpm), in acetate supplemented liquid medium (TAP) under LL (30 µmol photons m^-2^ s^-1^) or HL (500 µmol photons m^-2^ s^-1^), as indicated, or sparged with VLC in LL. Scale bar: 10 µm. **D.** Localization of a LCIB-mCherry fusion (magenta) monitored together with the position of the mitochondrial membranes (yellow) in cells grown mixotrophically as described in C. Fluorescence from the Chl channel was filtered out. Scale bar: 10 µm. **E.** Mitochondrial membrane locations in cells and their arrangement. Cells were layered on a poly-lysin-coated slide, topped with TP solid medium (1.5% low melting point agarose) and acclimated to HC or VLC conditions for 6 h. Dotted lines highlight the cells apico-basal axis as observed in cross sections. Scale bar: 10 µm. Fluorescence microscopy images are representative of 2 experiments. **F.** Cryo-EM tomogram of HC- or VLC-grown cells showing typical positions of mitochondrial membranes and distances between these membranes (mito) and the chloroplast (cp) or plasma membrane (pm). Red rectangles designate areas of very close proximity of the chloroplast envelope to the mitochondrial outer membrane. Scale bar: 200 nm.

When mitochondria are imaged as 3-dimensional reconstructions of whole cells (**Fig. 2E**), the tubular mitochondrial membranes appear as a network throughout the cell. Under HC conditions, this network resembles a web of highly reticulated, interconnecting membranes with no dominant orientation (**Fig. 2E**, **HC**). In contrast, under VLC conditions, the tubular membranes appear elongated over the surface of the chloroplast, oriented parallel to each other and often span the entire length of the cell, connecting at the poles (**Fig. 2E**, **VLC**). Quantification of the Clover fluorescence signal relative to the chloroplast position revealed a unimodal distribution of mitochondrial membranes at the cell periphery following VLC exposure, whereas in the HC-grown cells, most of the mitochondrial signal was within the inner cup of the chloroplast (**Supp. Fig. 1**). We also used cryo-electron tomography to determine the spatial relationships among mitochondrial membranes, the plasma membrane and the chloroplast envelop under HC and VLC conditions (**Fig. 2F**). In cells maintained under VLC conditions (**Fig. 2F, VLC)**, the mitochondrial membranes were often in close proximity to both the plastid outer envelope and the plasma membrane (**Fig. 2F, VLC**, red rectangle highlights close association between chloroplast and mitochondria). The distance between the outer chloroplast envelope membrane and mitochondrial outer membrane under VLC conditions is here less than 30 nm, extending over the envelope membrane surface of the plastid to lengths of several hundred nm (**Supp. Fig. 2**, video image).

The kinetics of mitochondrial relocation were analyzed after shifting cells from HC to VLC (**Supp. Fig. 3A** and **3B**, bottom). Mitochondrial relocation initiated ∼60 min after the shift and was mostly complete by 90 min (**Supp. Fig. 3A**). Upon a transition from VLC back to HC, the relocation of mitochondrial membranes to within the chloroplast cup appeared to take more than 120 min and was mostly complete after 180 min (**Supp. Fig. 3B**).

We conclude from these experiments that the Chlamydomonas mitochondrial network displays a massive rearrangement in response to VLC conditions, forming parallel, aligned tubules that extend in an apico-basal orientation at the cell periphery; this change in morphology takes about 90 min to complete and is readily reversible. Furthermore, mitochondrial relocation to the periphery appears to be the consequence of diminished intracellular CO_2_ levels, which is achieved by either decreasing the supply of Ci (TP, HC/LC to VLC) or by increasing internal CO_2_ consumption by photosynthesis (TAP, LL to HL), as suggested previously (17).

### Alignment of mitochondrial membranes requires microtubules

Mitochondrial motility and architecture in various organisms depend upon cytoskeleton components such as actin filaments, microtubules, and intermediate filaments (40), but photosynthetic organisms only harbor microtubules and actin. The Chlamydomonas genome contains genes encoding actin, NAP1 and IDA5, with the latter sensitive to the compound Latrunculin B (LatB) (41). Therefore, we examined the involvement of the actin cytoskeleton in mitochondrial relocation in the *nap1-1* mutant (41). While the absence of NAP1 in the *nap1-1* mutant had no effect on mitochondria relocation, treatment of *nap1-1* cells with LatB led to delayed relocation (**Fig. 3**). Whereas mitochondrial relocation and membrane reorganization were nearly complete in untreated *nap1-1* mutant cells after 90 min (**Fig. 3A**, DMSO 0.1%), a delay in the relocation was apparent in LatB treated *nap1*-*1* cells; even after 180 min, the cells still displayed a strong mitochondrial signal within the cup of the chloroplast (**Fig. 3A**). This experiment demonstrates that actin filaments are important for the timing of the mitochondrial relocation but are not required for the ultrastructural change of the mitochondria.

**Figure 3.**
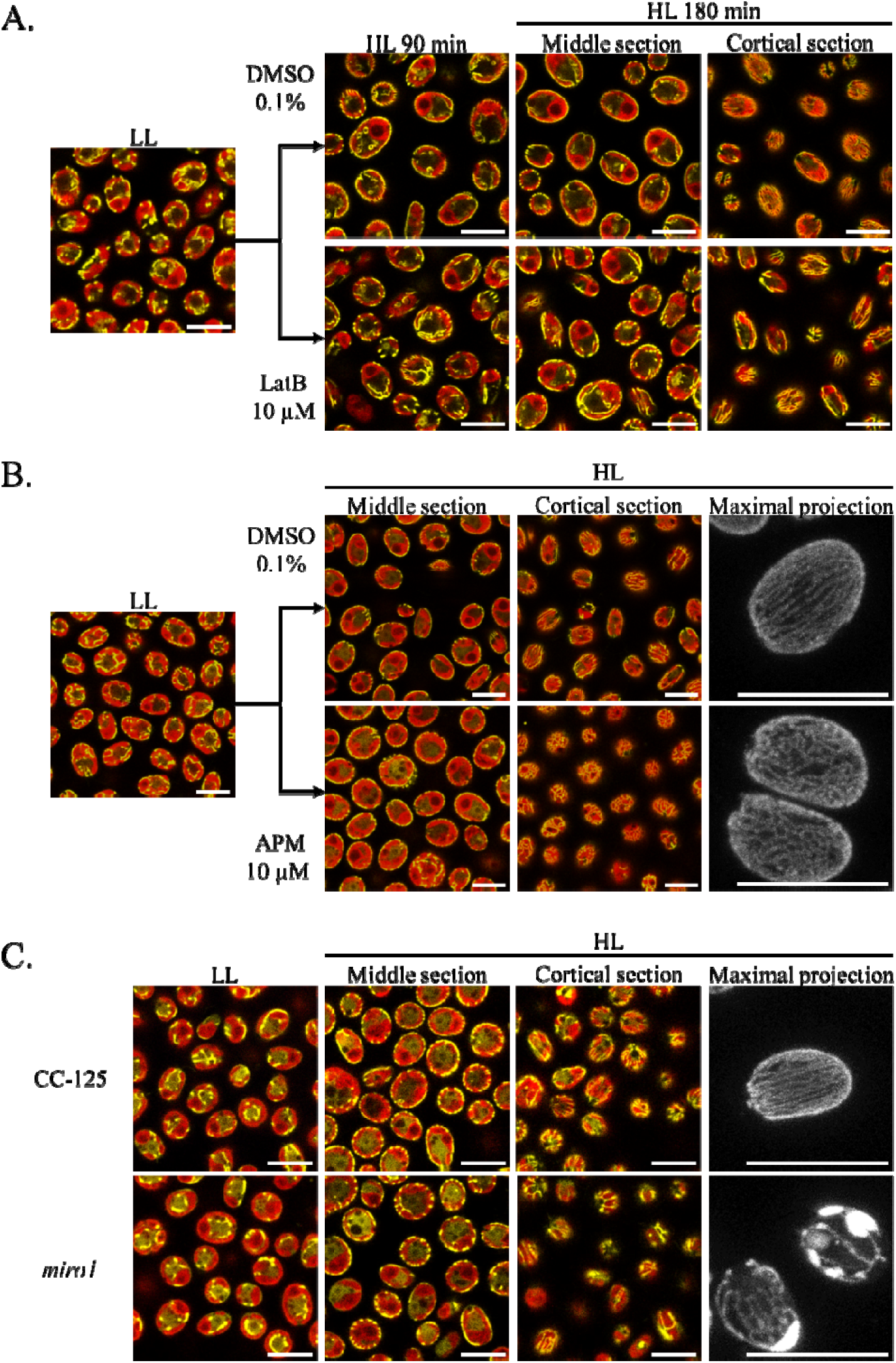
Involvement of cytoskeleton components on relocation of mitochondrial tubules. **A.** Effect of Latrunculin B (LatB) in the *nap1-1* mutant. Mitochondrial relocation induced by HL was examined in the *nap1-1* background, in the absence (only DMSO 0.1%) and presence (LatB 10 µM in DMSO 0.1%) of the actin inhibitor LatB. Scale bar: 10 µm. **B.** Effect of APM on mitochondrial location and membrane tubule organization. Mitochondrial relocation was examined in WT cells in the absence (only DMSO 0.1%) and presence of APM (APM 10 µM in DMSO 0.1%); cortical sections showing mitochondrial membrane organization near the plasma membrane and the maximal projections (cells immobilized on 1.5% TP agar) showing the whole cell mitochondrial signal. Scale bar: 10 µm. **C.** Effect of absence of MIRO1 upon HL induced relocation and the organization of mitochondrial membranes. Mixotrophically grown WT (CC-125) and mutant (*miro1*) cells were exposed to HL to induce mitochondria relocation. Cortical sections and maximal projections show the organization of the mitochondrial network. Scale bar: 10 µm. Fluorescence microscopy images are representative of 2 experiments.

The potential role of microtubules in the relocation process was assessed using the microtubule inhibitor Amiprophos Methyl (APM). Treatment with APM did not prevent the HL-triggered relocation of the mitochondrial membrane network to the cell periphery (**Fig. 3B**) but did disrupt the establishment of their apico-basal orientation; the mitochondrial tubules at the cell cortex appeared as an interconnected mesh of short tubules (**Fig. 3B**). Because the arrangement of mitochondrial tubules at the cortex extends from the apex to the base of the cell in a configuration similar to that of the cortical microtubules (42), we investigated colocalization of microtubules and mitochondria at the cortex using immunofluorescence. Mitochondria and microtubules were stained using specific antibodies directed against the mitochondrial CAH4 protein and α-tubulin, respectively (**Supp. Fig. 4**). Cortical microtubules spanned the entire cell length along the apico-basal axis (**Supp. Fig. 4, TUB**) and aligned with mitochondrial membranes when the cells were grown under VLC conditions; this alignment did not occur in HC (**Supp. Fig. 4, HC Merge**).

To further analyze the cytoskeleton requirement for relocation and reorientation of mitochondrial membranes, we investigated the effect of simultaneously eliminating actin and microtubules from the cells by simultaneously treating the *nap1-1* mutant with LatB and APM (**Supp. Fig. 5**). When both drugs are present, the effect is surprisingly severe relative to the loss of the individual cytoskeletal components (actin or microtubules). Mitochondria relocation is strongly inhibited during the first 90 min of exposure to HL (**Supp. Fig. 5, 90 min**). After 180 min, a fraction of mitochondria was at the periphery (**Supp. Fig. 5, 180 min**), but they appeared fragmented relative to control conditions, which might reveal whole cell defects that prevent attaining the physiological changes associated with VLC conditions.

In animal cells, mitochondrial interaction with microtubules is mediated by isoforms of MIRO, a conserved GTPase (43). The Chlamydomonas genome contains a single gene encoding a MIRO1 homolog (Cre08.g375200) (**Supp. Fig. 6**). To confirm the localization of MIRO1, we generated a strain expressing this GTPase fused with mCherry at its N-terminus. When expressed in VLC-grown cells, the mCherry-MIRO1 fusion aligned with the Clover signal, but also displayed increased intensity near the cell apex in the vicinity of the basal bodies, where cortical microtubules are organized (**Supp. Fig. 7A**). Mutants disrupted for *MIRO1* were generated by a CRISPR-guided insertion of a paromomycin resistance cassette in the background strain expressing the mitochondria-targeted Clover fluorophore (**Supp. Fig. 7B**). Independently isolated mutants had no defect on the relocation of mitochondria to the cell periphery but displayed significant structural aberrations in the mitochondrial membrane pattern under all growth conditions; there were large areas of aggregated/patchy mitochondria and generally lower numbers of mitochondrial membrane tubules that spanned the surface of the chloroplast under VLC conditions (**Fig. 3C, Supp. Fig. 7C**). These results indicate that microtubules and MIRO are required for attaining the parallel organization of mitochondrial membranes at the cell’s cortex under VLC conditions.

### Mitochondrial relocation correlates with CCM induction and is controlled by CIA5

Metabolic processes including photosynthetic electron transport, but also translation and transcription, could potentially be required for reorganization of mitochondrial membranes. To investigate the impact of photosynthesis on this reorganization, we used photosynthetic mutants and specific inhibitors during HL induction of mitochondrial relocation. Upon transfer of mixotrophically grown cells from VLL to HL, a mutant impaired in the primary reaction of photosynthesis (*F64* defective for the CP43 protein, does not accumulate PSII) (44) failed to induce mitochondrial relocation (**Fig. 4A, *F64***) while the control cells exhibited the expected pattern (**Fig. 4A, CC-124**). Similarly, a strain defective for regeneration of the Rubisco substrate ribulose biphosphate by phosphoribulokinase (PRK) (45) was also unable to induce mitochondrial relocation (**Fig. 4A, *prk***). The use of the PSII inhibitor DCMU or the PRK inhibitor Glycolaldehyde (GA) also inhibited relocation of mitochondrial tubules to the cell periphery (**Supp. Fig. 8A**). However, in the *F64* and *prk* mutant strains, the relocation of mitochondria was achieved when cultures where sparged with CO_2_-free air (**Fig. 4B**). In contrast, inhibition of mitochondrial respiration with Myxothiazol (MX, respiratory complex III inhibitor) did not block the relocation, although it did appear to result in fragmentation of the mitochondrial membrane network (**Supp. Fig. 8B**). We conclude that the cellular distribution of mitochondrial tubules depends primarily on the level of CO_2_ available to the cells, which is determined by (i) the rates of intracellular CO_2_ generation and consumption (which depends on respiratory and photosynthetic rates) and (ii) the Ci levels present in the environment.

**Figure 4.**
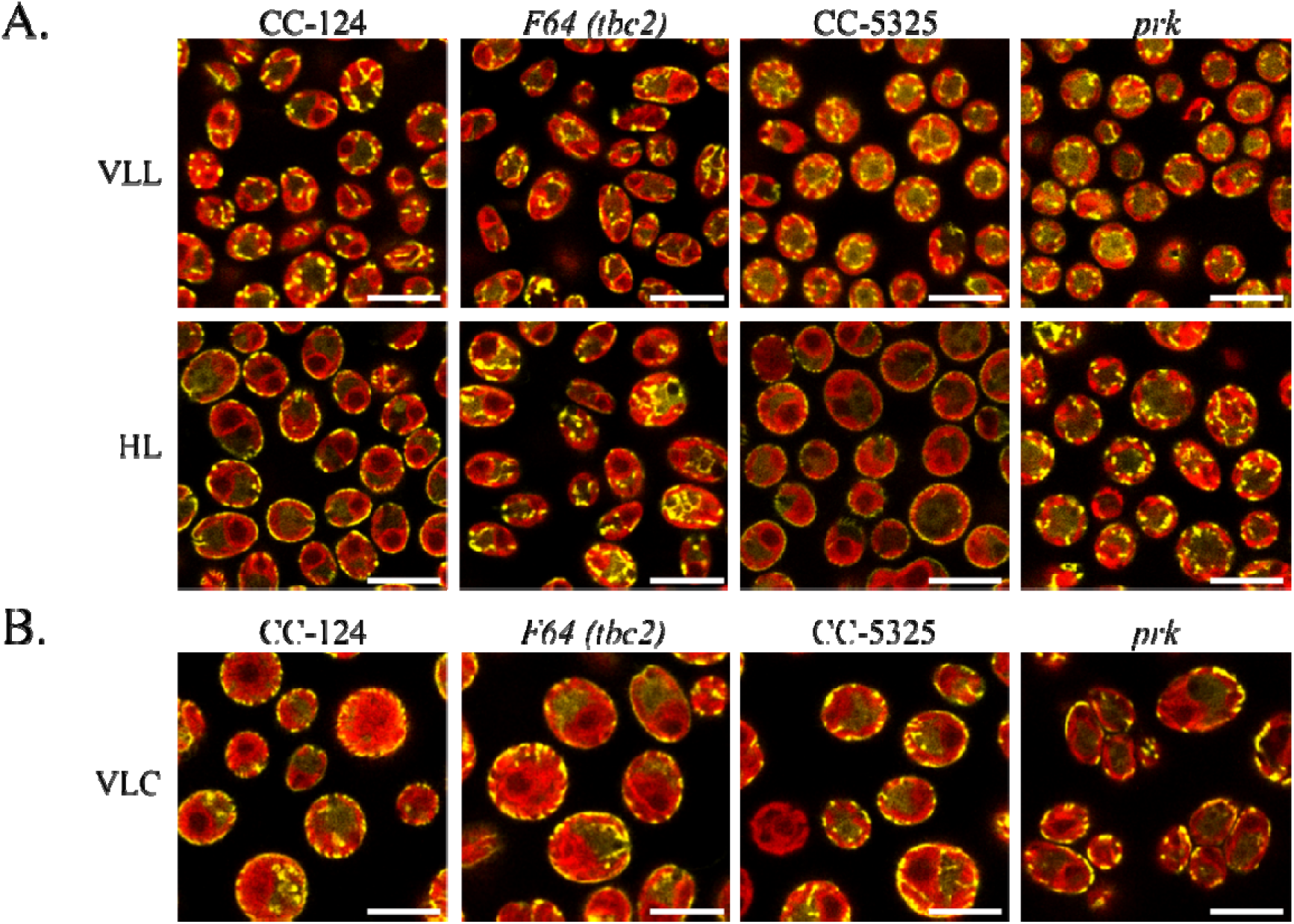
Effect of mutations that impact photosynthesis on mitochondrial relocation/reorganization. **A.** Effect of PSII and PRK mutations: mutant strains (*F64* and *prk*) and their corresponding parental strains (CC-124 and CC-5325) were mixotrophically grown in very low light (VLL, <5 µmol photons m^-2^ s^-1^) before being assayed for mitochondrial membrane rearrangement in HL. **B.** Effect of CO_2_ depletion on the position of mitochondria in the *F64* and *prk* mutants: mixotrophically grown cells were sparged with CO_2_ depleted air for 6 h and then assayed for their capacity for mitochondrial relocation under VLC conditions. Scale bar: 10 µm. Fluorescence microscopy images are representative of 2 experiments.

To assess the requirement for *de novo* gene expression and protein synthesis in driving mitochondrial rearrangements, we examined mitochondrial relocation in the presence of the eukaryotic transcription inhibitor actinomycin D (Act D), the eukaryotic translation inhibitor cycloheximide (CHX), and the chloroplast translation inhibitor chloramphenicol (CAP). Inhibitors of eukaryotic transcription and translation both prevented relocation of the mitochondrial membranes to the cell periphery (**Fig. 5A**) whereas the relocation was unaffected by the prokaryotic translation inhibitor CAP (**Supp. Fig. 9**).

**Figure 5.**
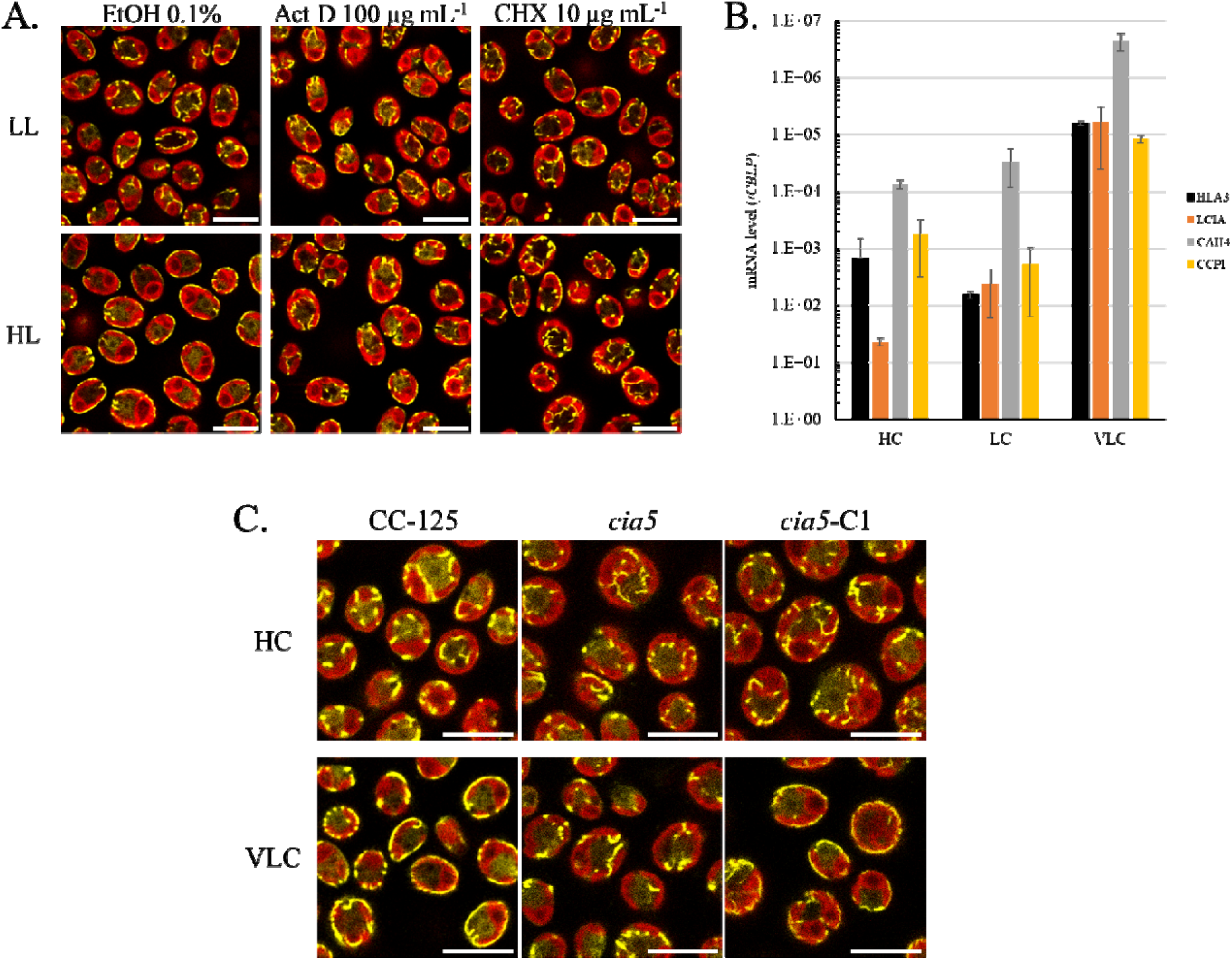
Relationship of mitochondrial relocation to the CCM. **A.** Effect of transcription and translation inhibitors. Cells were grown in TAP LL before HL treatment in the presence of ethanol (**EtOH 0.1%**), the transcription inhibitor actinomycin D (**Act D**), or the translation inhibitor cycloheximide (**CHX**). Control cells were incubated in LL in the presence of the drugs. Scale bar: 10 µm. **B.** Induction of CCM genes under conditions that cause movement of mitochondria to the cell periphery. The level of induction of genes encoding HLA3, LCIA, CAH4 and CCP1 in cultures sparged with HC (2% CO_2_), LC (air), and VLC (CO_2_-depleted air; <0.02%); all cultures were exposed to 100 µmol photons m^-2^ s^-1^. **C.** Dependence of mitochondrial relocation on CIA5. Wild-type (**CC-125**), the mutant (***cia5***), and the complemented (***cia5*-C1**) cells were grown photoautotrophically in HC and tested for mitochondrial relocation following 4 h of VLC treatment. Scale bar: 10 µm. Fluorescence microscopy images are representative of 2 experiments.

Because of the dependence of mitochondrial positioning on CO_2_ levels and *de novo* transcription/translation, we probed expression of CCM-related genes under the various CO_2_ conditions used to determine if there is a correlation between CCM induction and mitochondrial relocation. We grew the cultures under photoautotrophic conditions and transcript levels of known CCM genes were quantified by RT-qPCR following exposure of the cells to HC, LC and VLC (same conditions as in **Fig. 2A**). While transcript levels from the gene encoding LCIA increased steadily from HC to LC and VLC, transcripts levels of *HLA3* and the CCM associated mitochondrial genes, *CAH4* and *CCP1*, were only strongly induced when the cells were grown in VLC (**Fig. 5B**). The induction of mitochondria-localized proteins associated with the CCM paralleled mitochondrial relocalization to the cell periphery (**Fig. 2A**).

The correlation of mitochondrial relocation with the induction of CCM genes led us to test whether the former is governed by CIA5, the regulator that controls CO_2_-dependent expression of CCM genes (27, 28). We introduced the construct expressing the gene encoding a mitochondria-targeted Clover into the *cia5* mutant (32). VLC exposure of the *cia5* strain did not result in peripheral mitochondrial localization (**Fig. 5C**, ***cia5***) whereas this rearrangement was observed in the parental control strain (**Fig. 5C**, **CC-125**). Ectopic expression of a WT copy of the *CIA5* gene in the *cia5* mutant (**Supp. Fig. 10**) restored the strain’s ability for relocation (**Fig. 5C**, ***cia5*-C1**), demonstrating that CIA5 is integral to the relocation process. We also found that a mutant in the Ca^2+^ binding CAS1 protein, also linked to CCM gene expression (29), was not impacted for mitochondrial relocation (**Supp. Fig. 11**). We conclude that changes in the mitochondrial position and architecture upon acclimation of Chlamydomonas to VLC are controlled by the CCM regulator CIA5 and that they parallel changes in expression of the CCM-related genes.

### Mitochondrial relocation is correlated with an increased impact of respiration on CCM function

To further explore a potential role of changes in mitochondrial position/ultrastructure on CCM activity, we measured Ci dependent O_2_ evolution in WT cells to evaluate the apparent affinity of the cells for Ci under HC, LC and VLC conditions. Under LC conditions, CCM activation increases the cells affinity for Ci compared to HC, as shown by the reduced K_1/2_ value for Ci uptake (**Fig. 6A**), the concentration required to reach half maximum O_2_ evolution capacity. Growth under VLC conditions further increased the cell’s affinity for Ci (**Fig. 6A**).

**Figure 6:**
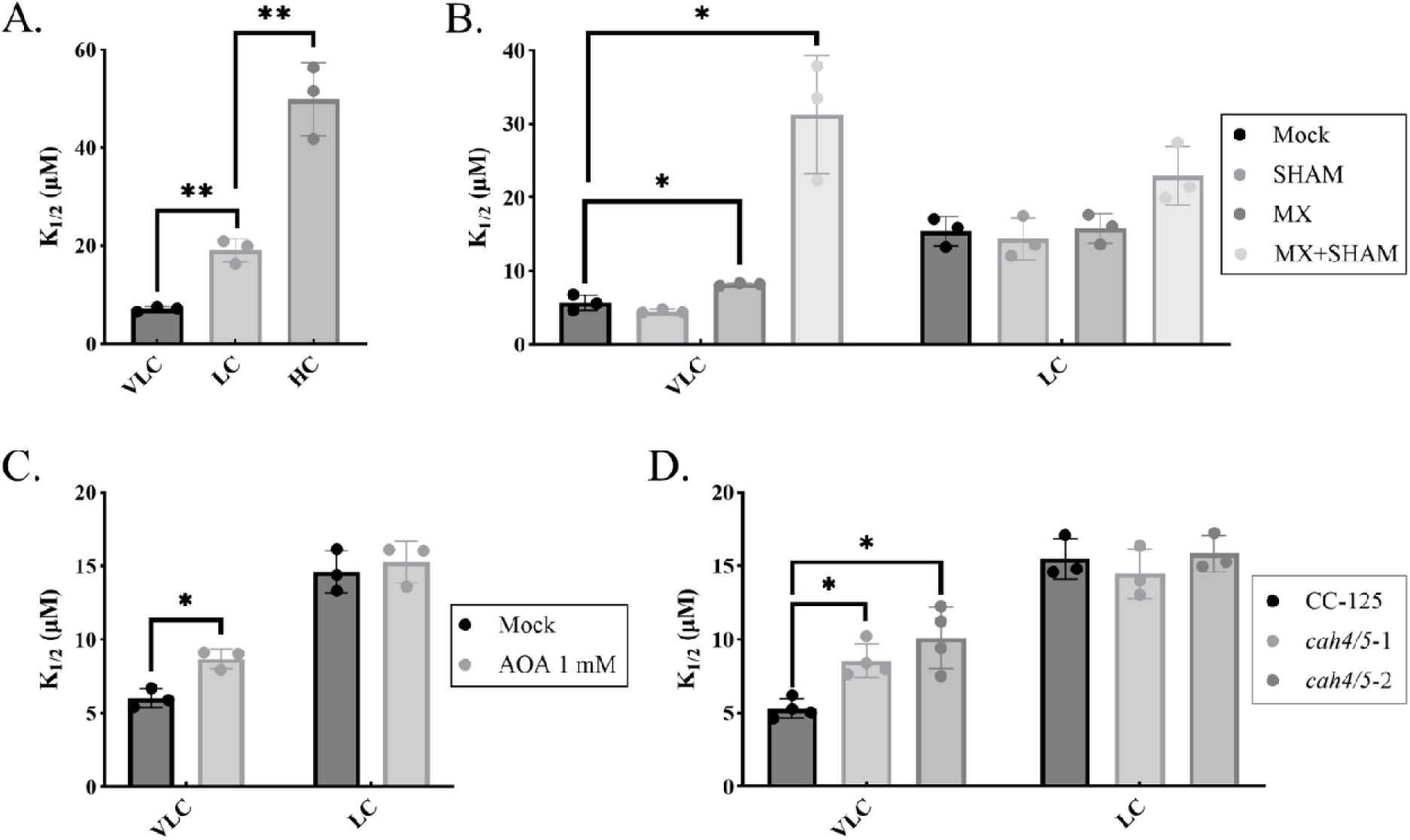
Analysis of mitochondrial inhibitors and mutants on Ci affinity. **A.** Ci affinity in WT cells grown in HC, LC and VLC conditions (n=3). **B.** Effect of the inhibitors SHAM and MX, separately and simultaneously on Ci affinity in cells grown in VLC and LC conditions. Inhibitors were added just prior to the assay (n=3). **C.** Effect of the aminotransferase inhibitor AOA (1 mM) on Ci affinity at VLC and LC. Cells were preincubated for 30 min in the presence of AOA (n=3). **D.** Ci affinity in the *cah4/5* mutants grown in LC and VLC conditions, compared to the WT parental strain (CC-125) (VLC, n=4; LC, n=3). * p value < 0.05. ** p value < 0.005.

To examine the potential role of mitochondrial electron transport on CCM activity, the cells were treated with the respiratory inhibitors MX and salicylhydroxamic acid (SHAM, alternative oxidase [AOX] inhibitor), as described in Burlacot *et al.* (37). While MX/SHAM treated VLC-grown cells showed a marked decrease in their apparent affinity for Ci (increased K_1/2_ value, **Fig. 6B**), LC-grown cells did not show a significant affinity difference upon MX/SHAM treatment (**Fig. 6B**). Each inhibitor was also used individually, with no effect for SHAM and a small but significant impact for MX under VLC conditions (**Fig. 6B**). We also tested cells disrupted for the apico-basal organization of the mitochondrial membrane network in both a mutant lacking the MIRO1 protein and WT cells treated with APM. Neither the mutant cells nor the APM treated WT cells showed a difference in their ability to grow under VLC conditions (**Supp. Fig. 7**) or their affinities for Ci relative to untreated WT cells (**Supp. Fig. 12A-B**).

Because it has been proposed to contribute to mitochondrial ATP production (38), we also tested the effect of inhibition of the photorespiratory pathway using the pyridoxal-phosphate analog aminooxyacetate (AOA), which inhibits transaminase reactions, including the reaction that converts glyoxylate to glycine during photorespiration, and results in the excretion of the accumulated intermediate glycolate (46) (**Supp. Fig. 12C**). We observed a decrease in the affinity of VLC grown cells for Ci after a 30 min exposure to AOA, but no effect was observed in LC (**Fig. 6D**).

We also tested the impact of the absence of two mitochondrial carbonic anhydrases, CAH4 and CAH5, known to be highly induced upon activation of the CCM and to be involved in CCM function (36). We generated knockout mutants for both *CAH4* and *CAH5* (**Supp. Fig. 13A**) and investigated their fitness and Ci affinity under VLC conditions versus LC conditions. We initially confirmed that the *cah4/5* double mutants were not impaired in their capacity to relocate mitochondria to the cell periphery (**Supp. Fig. 13C**). When incubated under LC conditions, the *cah4/5* mutants grew normally compared to the parental WT cells, but their growth was impaired under VLC conditions (**Supp. Fig. 13B**). Additionally, the mutants exhibited no difference in their affinity for Ci under LC conditions (**Fig. 6C, LC**) but had a lower affinity for Ci when grown in VLC (**Fig. 6C, VLC**). From these experiments we conclude that mitochondrial relocation in VLC is associated with an increase in the cell’s affinity for Ci, which occurs through the combined effects of respiratory activity, the activity of the mitochondrial CAH4/5 proteins and potentially photorespiration.

## DISCUSSION

Mitochondrial relocation during acclimation of Chlamydomonas to low CO_2_ was first observed by electron microscopy in the work of Geraghty and Spalding (15). Apart from significant vacuolization of the cells, probably a consequence of damaged chloroplast content and degradation of cellular components, the most striking change during acclimation of cells to low levels of Ci was the relocation of mitochondria to the cell periphery where they are wedged between the chloroplast outer envelope and the plasma membrane (15). Additionally, under H conditions, mitochondrial membranes were mostly in the cytoplasm within the cup formed by the chloroplast as a few, often large structures. Mitochondrial membrane vesicles from LC-grown cells appeared to be more numerous than those in HC grown cells; they also had a smaller cross-sectional diameter than those observed in HC grown cells, which was attributed to increased fragmentation or reticulation of the organelle as the level of CO_2_ declined. In this work, we investigated conditions and molecular factors required to induce and accomplish the peripheral positioning and changes in the structural organization of the mitochondrial network as the cells transition from HC to LC/VLC.

Relocation of the mitochondrial membrane network to the cell periphery was observed in photoautotrophic cultures that were experiencing VLC levels or when exposed to HL in acetate-containing medium. Mitochondria relocation was not observed when the cells were exposed to HC or LC. These results suggest that peripheral mitochondrial localization might be a feature of VLC acclimated cells (below ambient levels of CO_2_), which we supported by its correlation with the relocation of LCIB to the perimeter of the pyrenoid (VLC is required for LCIB relocation). Suppression of photosynthetic CO_2_ fixation, either using inhibitors or photosynthetic mutants, prevented HL-induced relocation, which occurred when the cultures were sparged with VLC. These results indicate that the signal triggering the rearrangements strongly depends on Ci availability in or around the cells, which is dictated by the balance between internal CO_2_ production, stimulated by acetate assimilation and respiration, photosynthetic CO_2_ fixation, which is elevated as the light levels increase, and the level of external Ci. Given the dependence of relocation/reorganization of mitochondria on Ci conditions and the production/consumption of CO_2_ by the cells, we investigated and confirmed the link between mitochondria relocation and CCM induction. Indeed, conditions that triggered mitochondrial redistribution were strongly associated with transcriptional induction of CCM genes, including *CCP1*, *CAH4, HLA3* and *LCIA*; the relocation was also dependent on CIA5, a protein critical for CCM induction. Together, our findings indicate that the Ci concentration is not only critical for controlling the CCM but is also a major factor that impacts the spatial distribution of the mitochondrial network in the cell. Whether the concentration of Ci directly or indirectly controls the cellular location of mitochondria, and the orientation of its membranes is not known.

Various hypotheses regarding a role for mitochondrial relocation to the cell periphery during VLC growth are depicted in **Fig. 7**. A recent study showed that the mitochondrial CAH4 and CAH5 proteins are required for optimal growth at low CO_2_ concentrations and contribute to Ci uptake (36). Our investigation of the *cah4/5* mutants showed a phenotypic defect even though the mitochondria achieved a peripheral location. Potentially, the CAHs could recycle the CO_2_ generated by mitochondrial respiration and photorespiration (*e.g.* routing Ci back to the chloroplast), but the peripheral mitochondrial position could also allow recapture of CO_2_ that might leak from the pyrenoid after HCO_3_^-^ is converted to CO_2_ by CAH3. In this case, the mitochondrion would form an additional barrier to CO_2_ leakage, augmenting the role associated with the starch sheath and the LCIB protein (putative CAH), both of which are positioned at the pyrenoid periphery under VLC conditions.

**Figure 7.**
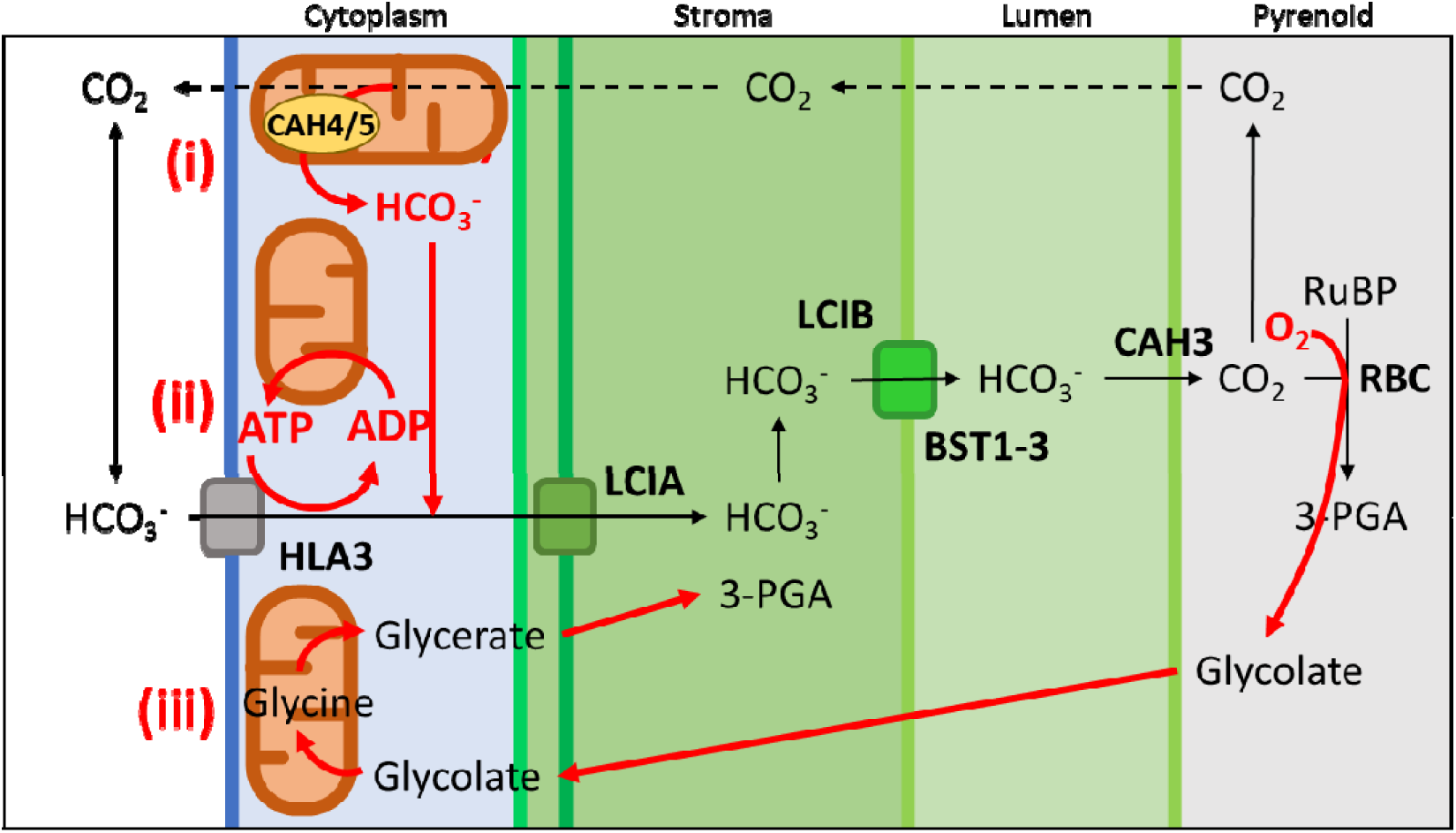
Possible roles of mitochondrial relocation to the cell cortex. (i) Mitochondrial CAH4/5 could capture CO_2_ that leaks from the chloroplast and channel it back into the chloroplast. (ii) Mitochondrial respiration can provide energy in the form of ATP for the active uptake of HCO_3_^-^ at the plasma membrane. (iii) Mitochondria can intercept glycolate and metabolize it through the photorespiratory pathway to limit the loss of Ci.

The *miro1* mutants displayed alterations in the alignment of the mitochondrial membrane network, although the relocation of the organelle to the cell periphery was normal under VLC conditions. Additionally, CCM induction and function in response to VLC were not impaired in the mutants despite a clear reduction in the coverage of the chloroplast surface by the mitochondrial membranes. Instead of having regularly spaced, parallel membrane tubules spanning the length of the cells, *miro1* cells had a reduced number of mitochondrial membrane tubules, leaving large areas of the chloroplast surface devoid of mitochondrial membranes. These findings suggest that the principal function of mitochondrial relocation is unlikely to be that of limiting CO_2_ leakage from the cell. CAH4/5 accumulation could potentially salvage CO_2_ produced by respiration and photorespiration, support the Krebs cycle by maintaining the supply of HCO_3_^-^ needed for oxaloacetate replenishment (47), or both. Supplying metabolites for the Krebs cycle would ensure continued production of NADH for respiration and ATP synthesis.

Additionally, the peripheral mitochondrial position could serve as a physical barrier to limit glycolate leakage from cells. Although expected to be a minor metabolite once the CCM becomes fully active (48), photorespiratory generated glycolate is exported by the chloroplast and could also leak from cells; this glycolate might be more efficiently recycled by the cortex-localized mitochondria (46). The photorespiratory pathway may also integrate with CAH4/5 function to facilitate CO_2_ recycling. However, we would expect this process to be impaired in the *miro1* mutants and exhibit a CCM phenotype. Since the mutants do not appear to be CCM compromised, the recycling of glycolate would, at most, be a minor aspect of mitochondrial relocation.

Mitochondria may also contribute to CCM function by providing ATP to energize HCO_3_^-^ transporters (HLA3) on the plasma membrane (37). Spatially optimizing the site of ATP supply and utilization has been observed in neuronal cells and during development (3, 4, 49). The peripheral mitochondrial localization could physically optimize the use of energy resulting from colocalization of respiratory ATP production and consumption by plasma membrane Ci transporters. We currently favor this hypothesis since inhibition of complex III by MX impacts the cell’s affinity for Ci under VLC conditions while SHAM had no apparent effect. Indeed, the complex III pathway promotes translocation of two more protons than the AOX pathway and its inhibition has a more severe impact on ATP production than does the inhibition of AOX. This hypothesis is congruent with the finding that mitochondrial relocation is a VLC response since, under VLC conditions and according to Wang and Spalding (21), the cells mostly rely on HCO_3_^-^ uptake from the environment, which would require plasma membrane and chloroplast envelope HCO_3_^-^ transport activity. Under such conditions, HLA3 activity can be supported by mitochondrial respiration while the LCIA channel would mediate passive HCO_3_^-^ diffusion (23). Additionally, the plasma membrane putative H^+^ exporting ATPase ACA4, which is proposed to aid HCO_3_^-^ uptake (50), may also benefit from the mitochondrial ATP supply.

Participation of mitochondria in CCM function in other algae has also been observed. In diatoms, mitochondria are strikingly different from those of Chlamydomonas; they are much less reticulated and more globular, which may impact their potential for major rearrangements (51). However, there is evidence of physical interactions of mitochondria and chloroplasts that may be modulated by growth conditions, possibly promoting the exchange of metabolites between these organelles. In *Nannochloropsis oceanica* (family Eustigmataceae) and in the diatom *Phaeodactylum tricornutum*, recent studies have suggested the importance of the mitochondrial phosphoenolpyruvate carboxylase for optimal growth when Ci becomes scarce, likely through HCO_3_^-^ acquisition in the form of oxaloacetate (52, 53). Despite its partial use of an apparent C4-type biochemical CCM, *N*. *oceanica* also relies on a biophysical CCM in which LC-induced, mitochondrial CAHs could be involved, as indicated by the lower fitness of specific RNAi strains under LC conditions (54). *Thalassiosira pseudonana* also maintains mitochondria-targeted CAH activity (55). While the structural dynamics of mitochondria in response to Ci limitations have not been broadly studied in microalgae, rearrangements of mitochondria similar to those observed in Chlamydomonas have been noted in *Scenedesmus obliquus* (16) and suggested for *Chlorella ohadii* (56) and *Volvox africanus* (57). Intriguingly, a peripheral mitochondrial position has also been observed in non-photosynthetic relatives of Chlamydomonas (58–61), so it seems likely that the peripheral localization captures some function(s) not exclusively associated with supporting photosynthesis.

While VLC induced relocation is readily observed by fluorescence microscopy, it is reasonable to assume that the subcellular localization of mitochondria can be tailored to locally provide energy/ATP to various cellular processes. In Chlamydomonas, a portion of mitochondrial tubules is consistently observed around the contractile vacuoles and their movement tracks the rhythmic beating of the vacuole (**Supp. Fig. 14**), likely providing ATP for vacuolar proton pumping ATPases. The close association between mitochondria and chloroplasts could also optimize the use of mitochondrial generated energy for the import of proteins through the TIC-TOC complex, which is mainly localized near the plastid lobes (50).

The movement and positioning of mitochondria occur in many organisms in addition to algae and has been shown to be dependent on the actin and microtubule cytoskeleton proteins (40); this phenomenon remains poorly understood in plant cells, and even more so in algae like Chlamydomonas. The striking parallel organization of mitochondrial membrane tubules is dependent on the microtubules, to which the mitochondrion probably attaches. However, in contrast to the parallel arrangement of the mitochondrial membranes, the peripheral mitochondrial localization was not affected by treating cells with microtubule inhibitors. Additionally, there was no marked requirement for the actin cytoskeleton during VLC-triggered repositioning of mitochondria, although it might speed-up the relocation process. Simultaneous inhibition of microtubule and actin cytoskeleton formation strongly impacted relocation, although dramatic fragmentation of the mitochondrial network in these cytoskeleton deficient mutants may indirectly impact the internal cellular architecture and suppress mitochondrial membrane relocation. An association between microtubules and mitochondria at the cell’s cortex under VLC conditions may be mediated by adaptor proteins that link microtubules to the outer mitochondrial membrane. One mediator of mitochondrial-cytoskeletal interaction is MIRO1, a protein critical for mitochondrial network dynamics in mammals, yeast, and plants (62–64). Inactivation of the *MIRO1* gene dramatically impacts the parallel arrangement of mitochondrial membranes but the mitochondria still relocated to the cell periphery upon exposure to VLC and do not appear to be impacted in their efficiency of Ci utilization. While the impact of MIRO1 loss on the mitochondrial membrane arrangement might result from the absence of microtubule/mitochondria interactions, it may also be a consequence of disruption of the ERMES complex (ER-mitochondria encounter structure), identified in yeast and mammals (65), which controls the fission of mitochondria and mediates mitochondria-ER lipid exchange through contact sites between the compartments; MIRO1 is a known interactant/component of this complex (66, 67). A more in-depth investigation of mitochondrial movement will be necessary to identify those molecular players involved in repositioning the mitochondrial to the cell periphery.

Despite the striking correlation between mitochondrial organization and its spatial location in the cells upon induction of the CCM, it is not clear how important this structural flexibility is for growth under VLC conditions. Therefore, elucidating the molecular mechanisms associated with mitochondrial movement might not be amenable to a mutant screen for altered growth under VLC conditions. One promising strategy would involve screening for mutants unable to perform VLC-induced mitochondrial relocation based on single cell imaging. To this end, intelligent image assisted cell sorting (IACS) (68) is being used to develop high throughput screening of insertional mutants. This system has been used for screening Chlamydomonas transformants that are unable to concentrate LCIB around the pyrenoid when the cells are shifted to VLC (68). Our preliminary results have demonstrated the feasibility of enriching for ‘relocation’ mutants based on the movement of mitochondria to the cell periphery under VLC conditions (39). This technology will help elucidate the importance of mitochondrial relocation under VLC conditions and identify genes/proteins required for this redistribution.

## METHODS

### Strains

CC-125 and the *cia5* mutant (32) were provided by Dr. Dimitris Petroutsos (CEA, Grenoble, France). The photosynthetic mutant F64 was a gift from the Institut de Biologie Physico-Chimique (IBPC, Paris, France) (44); it was compared to its parental strain, CC-124. The *prk* mutant was a gift from Dr. Pierre Crozet (Sorbonne Université, Paris, France) (45) and the *cas1* mutant was ordered from the CLiP library (LMJ.RY0402.131739) (69). The parental strain for both mutants is CC-5325. The actin mutant *nap1-1* was a gift from Dr. Masayuki Onishi (Duke University, Durham, USA) (41). The WT strain WT222+ was used for generating cryo-ET images. Strain CSI_FC1D06 constitutively expresses the mitochondrial localized CAH4 protein (Cre05.g248400) fused to the Venus fluorophore (50). This construct has been useful to view mitochondria by immunofluorescence in HC and VLC. All newly generated strains are available at the Chlamydomonas Resource Center (https://www.chlamycollection.org/).

### Plasmid construction

The plasmid expressing a mitochondria-targeted Clover fluorophore was constructed from components of the Modular Cloning kit for Chlamydomonas (70). The Clover gene (B3-B5) was fused to the sequence encoding the mitochondrial matrix HSP70C targeting peptide (B2) and the fusion protein was expressed under the control of the pAR promoter (A1-B1, HSP70A/RBCS2 fusion promoter) (71) and the PSAD terminator (B6-C1) (72). This construct was assembled in a pLM1 level plasmid harboring the hygromycin resistance cassette unless the target strain was already hygromycin resistant, in which case paromomycin resistance was used for selecting transformants. The *LCIB* gene was cloned into the pLM006 backbone (50) in frame with the mCherry fluorophore sequence using primers listed in **Table 1**.

**Table 1:**
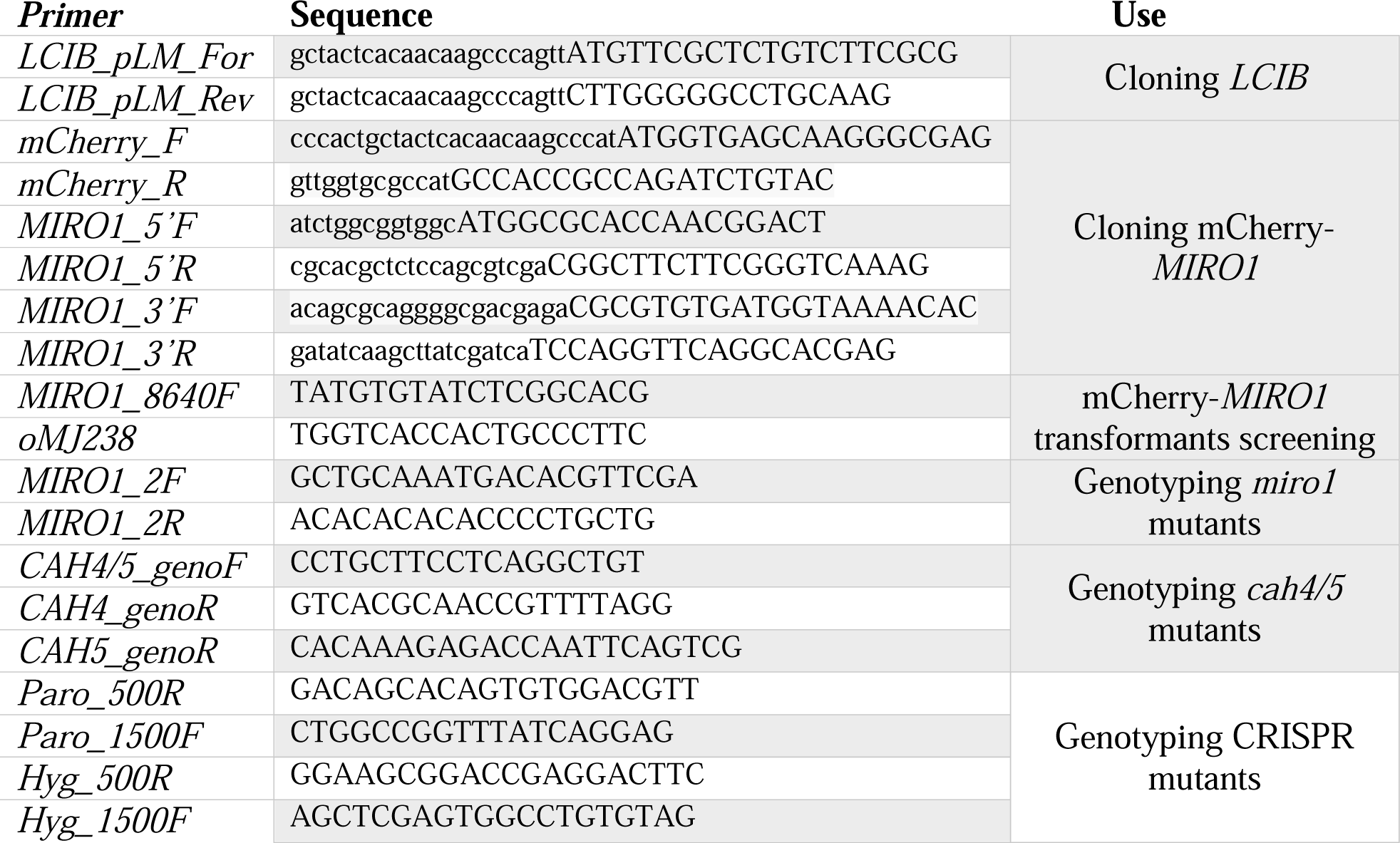
PCR primers.

The *MIRO1* gene was cloned into the pSLSpec backbone (pSL18 modified to encode spectinomycin resistance) in an N-terminal fusion (MIROs are tail anchored proteins) with mCherry that was generated by Gibson assembly (New England Biolabs, Ipswich, MA, USA; see primers in **Table 1**). The pSLSpec plasmid was linearized with NdeI and 3 segments of the *MIRO1* gene (5’ 479 bp from start codon, 8137 bp fragment from PTQ2712 SalI/MluI restriction, 3’ 647 bp until end of 3’UTR) were inserted into the vector just downstream of the mCherry coding sequence that was amplified from pLM006 (50). pSLSpec-mCherry-MIRO1 was linearized and transformed into the *miro1* mutant.

### Culture conditions

Cells were maintained on TAP agar medium for long term storage. Appropriate antibiotics were included in the solid medium to avoid the loss of expression of the introduced fluorophore. Batch cultures were grown in liquid TAP medium at low light (LL, 30 µmol photons m^-2^ s^-1^) or very low light (VLL, <5 µmol photons m^-2^ s^-1^), and the cells were diluted and allowed to grow overnight to ensure logarithmic growth at the time of the assay.

For determining the distribution of mitochondrial membranes under different light intensities and in the presence of different carbon sources, cultures were inoculated at 1 µg Chl mL^-1^ in either 30 mL of medium in 125 mL Erlenmeyer flasks that were stoppered with aluminum foil, or in 50 mL of medium in 2.5 cm wide, 20 cm long glass tubes with aeration from the bottom provided by a Pasteur pipette. The cells were grown for 24 h under various conditions, as specified in the text. VLC conditions were created by bubbling air twice through 50% sodium hydroxide, which reduced the CO_2_ level in the medium to ∼150 ppm as determined by measurements with an Amprobe CO_2_ meter (Everett, WA, USA).

For imaging the mitochondrial membrane distribution, the cells were concentrated and layered on a poly-lysine-coated 8-chamber slide (Ibidi, Martinsried, Germany) and topped with 300 µL of 1.5% low-melting point agarose in TP/MOPS medium at 34°C. Cells were incubated at 100 µmol photons m^-2^ s^-1^ under HC or VLC atmosphere for 6 h to allow for recovery from mild temperature shock and to stabilize the mitochondrial membrane distribution. For monitoring relocation kinetics, the cells were layered on a slide in TP medium as described. Redistribution of mitochondrial membranes was induced by exposure of HC-maintained cells to ambient conditions (ambient levels of CO_2_ at 100 µmol photons m^-2^ s^-1^). After stabilization of the peripheral mitochondrial distribution (6 h minimum), the light was turned off to allow the mitochondrion to revert back to a more internal (central in the cell) mitochondrial membrane distribution.

When testing drugs and specific mutants, induction of mitochondrial membrane relocation was performed under mixotrophic conditions. Cells were grown in TAP medium at LL (or VLL for photosynthetic mutants) and a 5 mL aliquot was exposed to HL for 3 h. GA (5 mM), DCMU (10 µM), MX (2.5 µM), SHAM (400 µM), CHX (10 µg mL^-1^), Actinomycin D (10 µg mL^-1^), AOA (1 mM) and APM (10 µM) were purchased from Sigma-Millipore (Saint-Louis, MO, USA), and Latrunculin B (10 µM) from abcam (Cambridge, UK). A 1 mL aliquot of cells at 1-2 × 10^6^ cells ml^-1^ was concentrated 20-fold after pelleting the cells by centrifugation for 1 min at 1500 x *g.* The concentrated suspension was deposited on a poly-lysine-coated 18-well slide (Ibidi, Martinsried, Germany).

For cryo-electron tomography, WT222+ cells were grown to mid-log phase in TAP medium, pelleted by centrifugation, washed once with High Salt Medium (HSM), resuspended in a small volume of HSM and diluted in TAP or HSM to an OD_750 nm_ of 0.1. The cells were then allowed to acclimate to the fresh medium in the dark (TAP) or at 60 µmol photons m^-2^ s^-1^ (HSM) for 24 h.

### Transformation and screening

Exponentially growing cells were transformed using the GeneArt Max Efficiency Reagent (Thermo Fisher, Waltham, MA, USA) following the manufacturer’s instructions. Transformants were selected with appropriate antibiotic(s) (hygromycin B, 20 µg mL^-1^; paromomycin, 10 µg mL^-1^; spectinomycin, 100 µg mL^-1^ Sigma-Millipore) for a week before screening them for fluorescence in 96 well plates. Resistant colonies were picked, arrayed on solid medium and the toothpick used for placing cells on the solid medium was also dipped into 100 µL of TAP medium for direct inoculation of the 96 well plates. After two days of growth in LL with agitation, the plates were scanned with a Tecan M1000 plate reader. Clover fluorescence was monitored at 500-530 nm after excitation at 488 nm. mCherry fluorescence was monitored from 570 to 610 nm after excitation at 561 nm. Every clone that we selected displayed a fluorescence value that was at least 3 times above background (background values were from the non-transformed parental strain) and then validated by confocal microscopy.

### Confocal microscopy

A TCS SP8 confocal laser-scanning microscope (Leica) with imaging conditions/settings as follows: cells were imaged using LASX software at a ×63, numerical aperture and a 1.4 oil objective. Excitation/emission settings were 514 nm (notch filter)/525–550 nm HyD1 SMD hybrid detector for Clover, and 514 nm/680–720 nm HyD2 SMD hybrid detector for chlorophyll autofluorescence, working in parallel. The EM gain was set at 100%. Clover fluorescence was captured with a lifetime gate filter (0.6-10 ns) to reduce background noise from chlorophyll autofluorescence. Z-stacks were collected at 0.2 μm intervals to generate maximum projections. Images were analyzed using Fiji software.

### Quantification of the Clover fluorescence signal

A custom algorithm was assembled in LabView and used to quantify the radial distribution of a mitochondria-associated fluorescence signal. Briefly, individual cells were cropped from 3D data, the ellipsoids were then rotated by 2 angles in 3D space and fitted to the overall chloroplast autofluorescence signal. Relative mitochondrial and chloroplast signals in ellipsoids of proportionally decreasing sizes were normalized to the total cumulative signal of the cells. The method will be described in more detail elsewhere (manuscript in preparation).

### Cryo-electron Tomography

After 24 h of culturing Chlamydomonas cells either in VLC (HSM, 60 µmol photons m^-2^ s^-1^) or HC (TAP, Dark), specimens were vitrified by plunge-freezing 3 µL aliquots of the cell suspension onto glow-discharged Quantifoil Multi A Holey Carbon Au 200 mesh TEM grids (SPI Supplies, USA) using a Leica EM GP2 apparatus (Leica, Austria), in which the humidity was kept at under 95% with 4 to 8 seconds of single-sided blotting on the reverse side of the grid. Ultrathin lamellae (150-200 nm) were prepared using an Aquilos2™ cryogenic Focused Ion Beam Scanning Electron Microscope (cryoFIB-SEM, ThermoScientific, USA), operated using a 2-5 kV electron beam, a 30 kV Ion beam with the ion probe current adjusted from 0.3 nA to 30 pA for rough milling to final polishing. Grids were transferred under cryogenic conditions to a 300 kV Krios™ cryogenic Transmission Electron Microscope (cryo-TEM, ThermoScientific, USA) with a Gatan K3 detector and BioQuantum energy filter (Gatan Inc, USA) for cryo-ET data collection, which used a pixel size of 3.4 Å, a -60 to 60 degrees tilt range at 2-degree increments, and Serial EM software for automated data collection (73). Tomograms were generated using IMOD software (74) followed by segmentation and visualization using EMAN2 (75) and USCF Chimera (76).

### Immunofluorescence

VLC- and HC-grown cells were harvested by centrifugation for 3 min at 1500 x *g*. The pellets were resuspended in MT buffer (77) containing 3.3% paraformaldehyde. Cells were incubated 5 min at room temperature, pelleted by centrifugation, resuspended in MT buffer containing 0.5% NP-40 for 2 min and then transferred to MT buffer and layered onto a poly-lysine coated 8-chamber slide for 15 min to allow attachment. The buffer was then removed, and the cells allowed to air dry for 5 min. The slide was immersed in -20°C cold methanol for 10 min (in a Coplin Jar) to remove the chlorophyll, followed by rehydration by 3 successive, 10 min incubations in PBS. These cell samples were then incubated in PBS + 5% BSA for 60 min at room temperature for blocking (elimination of unspecific signal). The blocking solution was replaced by PBS-T (0.1% Tween-20) containing primary antibodies, anti-CAH4/5 (1/500 dilution; AS11 1737, Agrisera, Vännäs, Sweden) and anti-α-Tubulin (B-5-1-2, 1/500 dilution; Santa Cruz Biotechnology, Dallas, TX, USA). After a 4 h incubation with the primary antibodies at 37°C, the slide was washed 3 times for 5 min with PBS followed by the addition of secondary antibodies to the PBS-T (Alexa Fluor 488 nm, anti-rabbit, 1/500 dilution; Alexa Fluor 568 nm, anti-mouse, 1/500 dilution; Invitrogen, Waltham, MA, USA) and incubated at room temperature for 1 h in the dark. The cells were then washed 3 times for 5 min with PBS and covered with PBS before imaging them with a TCS SP8 confocal laser-scanning microscope (Leica). Excitation/emission settings were 488 nm/498-524 nm HyD1 SMD hybrid detector for CAH4, and 580 nm/627-672 nm HyD2 SMD hybrid detector for α-Tubulin.

### CRISPR/Cas9 mutagenesis

The *miro1* and *cah4/5* mutants were generated according to Findinier *et al.* (78) with some modifications. Prior to transformation, the cell wall was digested by incubating the cells in an autolysin solution for 45 min at 100 µmol photons m^-2^ s^-1^ without shaking (79). Cells were then washed twice with 50 mL of TAP 40 mM sucrose medium. During cell wall digestion, the ribonucleoprotein (RNP) complex was prepared with Cas9 protein (IDT, Coralville, IA, USA) and sgRNA targeted to the *MIRO1* gene (5’-CTCGTCCAAGTTCAACAAGATGG-3’; exon 3, sense) or to the *CAH4* and *CAH5* genes (5’-GCCCTGGAGTACCTTCGCGAGGG-3’; exon 2, sense), and incubated at 37°C for 30 min. WT cells were co-transformed with the RNP complex and a resistance cassette (*aphVIII* amplified from pSL18 for *MIRO1*, *aph7”* amplified from pSLHyg as in Findinier et al. (78) for *CAH4/5*; see **Table 1**). After selection on antibiotic containing agar medium, transformants were screened by PCR using a primer in the 5’ or 3’ region of the resistance cassette, and oriented outward coupled with a pair of primers around the target site on the specific gene (**Table 1**).

### Transcript quantification

HC, LC and VLC grown cells were pelleted for 2 min at 800 x *g*, the supernatant discarded, and the pellet flash frozen in liquid nitrogen. Total RNA was extracted using the Qiagen RNA easy extraction kit following the manufacturer’s instructions. cDNA was synthesized from total RNA using the iScript Reverse Transcription Supermix (Bio-Rad, Hercules, CA, USA) with 1 µg of RNA in a 20 µL reaction volume. cDNAs were diluted to 50 µL and used as template for Real-Time PCR monitored by SensiFast SYBR No-Rox Kit (Bioline, Cincinatti, OH, USA). RT-PCR was performed in the Roche Light Cycler 480 using primers for *CAH4*, *CCP1*, *LCIA*, *HLA3* and *CBLP* (**Table 2).**

**Table 2:**
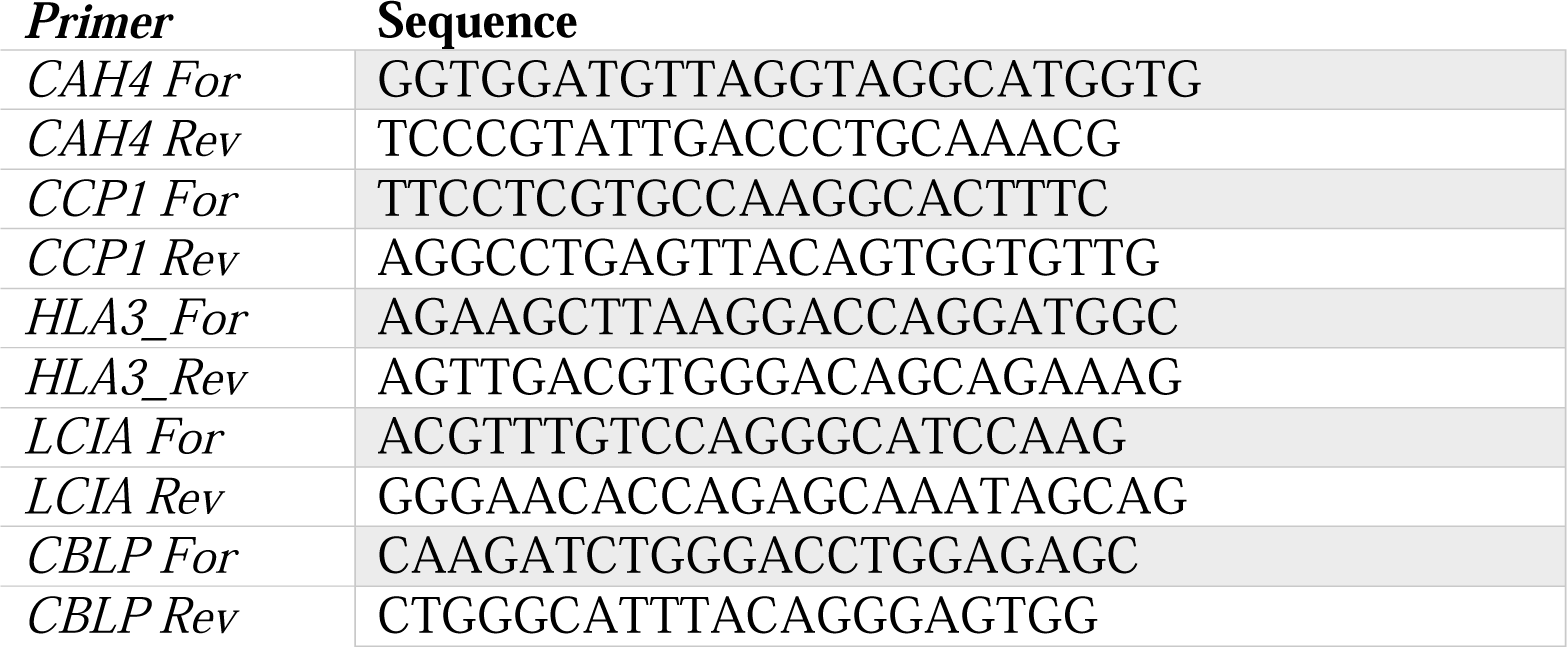
RT-qPCR primers.

### Complementation

*cia5* and *cas1* complementation took advantage of the ability of the complemented strains to grow under VLC conditions. A WT copy of the *CIA5* gene was excised from the BAC plasmid PTQ9468 by MluI restriction (7438 bp fragment) and transformed into the *cia5* mutant expressing the mitochondria-targeted Clover fluorophore. The *CAS1* gene was excised from PTQ9603 (5656 bp) with the restriction enzymes KpnI/NruI and transformed into the *cas1* mutant harboring the Clover transgene. For both mutants, control transformations were performed using the plasmid DNA restricted by an enzyme that cuts the target gene within its coding sequence. No strains that grew at VLC were obtained from the control transformations.

### Oxygen evolution measurements

HC, LC and VLC-grown cells were pelleted for 3 min at 1,800 x *g* and resuspended at 20 µg Chl mL^-1^ in 2 mL of fresh medium that was sparged with CO_2_-free air. The cells were then loaded into the sample chamber of an Oxygraph+ oxygen electrode system (Hansatech, Norfolk, England). The chamber was sealed and the cells were exposed to 300 µmol photons m^-2^ s^-1^. Once the O_2_ evolution rate declined to net zero, NaHCO_3_ was added to the suspension at a final concentration of 10, 50, 250 and 1250 µM and the O_2_ evolution rate measured. The K_1/2_(Ci) was calculated from the fitted curve for each strain. AOA was added at 1 mM to the cell culture 30 min prior to measurements of O_2_ evolution. MX and SHAM were added at 2.5 µM and 400 µM, respectively, during the CO_2_ depletion period.

### Glycolate measurements

Cells were grown for 24 h in VLC in the absence or presence of the inhibitor AOA (1 mM). Cells were then centrifuged for 5 min at 1,500 *g*, and 1 mL of supernatant was harvested and freeze-dried. Glycolate was measured according to Takahashi (80).

## ACKNOWLEDGEMENT

This project was supported by funding from DOE award DE-SC0019417 (to A.R.G) and BERFWP 100463 (to W.C.). JF was supported by the Department of Plant Biology of the Carnegie Institution for Science. Some of this work was performed at the Stanford-SLAC CryoET Specimen Preparation Center (SCSC), which is supported by the National Institutes of Health Common Fund’s Transformative High Resolution Cryo-electron Microscopy program (U24GM139166). We would like to thank the Carnegie Advanced Imaging Facility for use of microscopy instruments (Leica TCS SP8 confocal laser-scanning microscope).

