## Supplemental figures and tables for "Dramatic Changes in Mitochondrial Subcellular Location and Morphology Accompany Activation of the CO_2_ Concentrating Mechanism"

**Supplemental Figure 1.** Quantification of the distribution of mitochondrial fluorescence signal relative to an ellipsoid shaped cell defined by chlorophyll autofluorescence. Cells incubated in HC and VLC were analyzed. (n=15). Open circle: distribution of chlorophyll signal in HC; closed circle: distribution of chlorophyll signal in VLC; open square: distribution of mitochondrial Clover signal in HC; closed square: distribution of mitochondrial Clover signal in VLC.

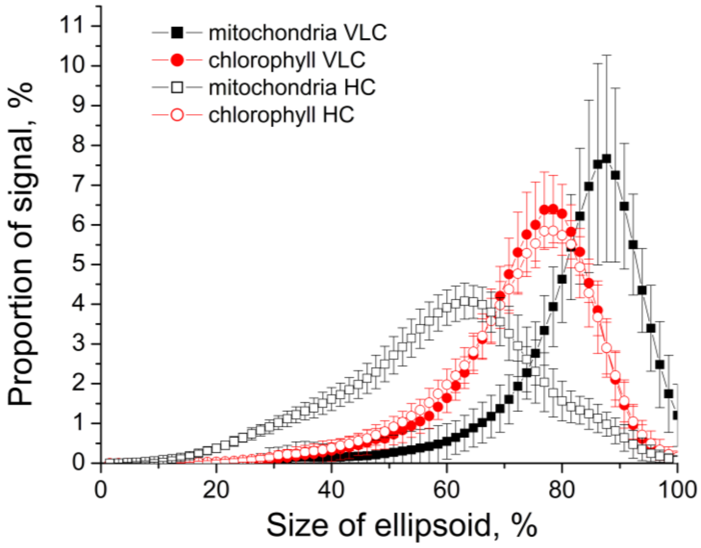

**Supplemental Figure 2**. Images and videos of the tomogram generated from HC and VLC conditions. Scale bar: 200 nm.

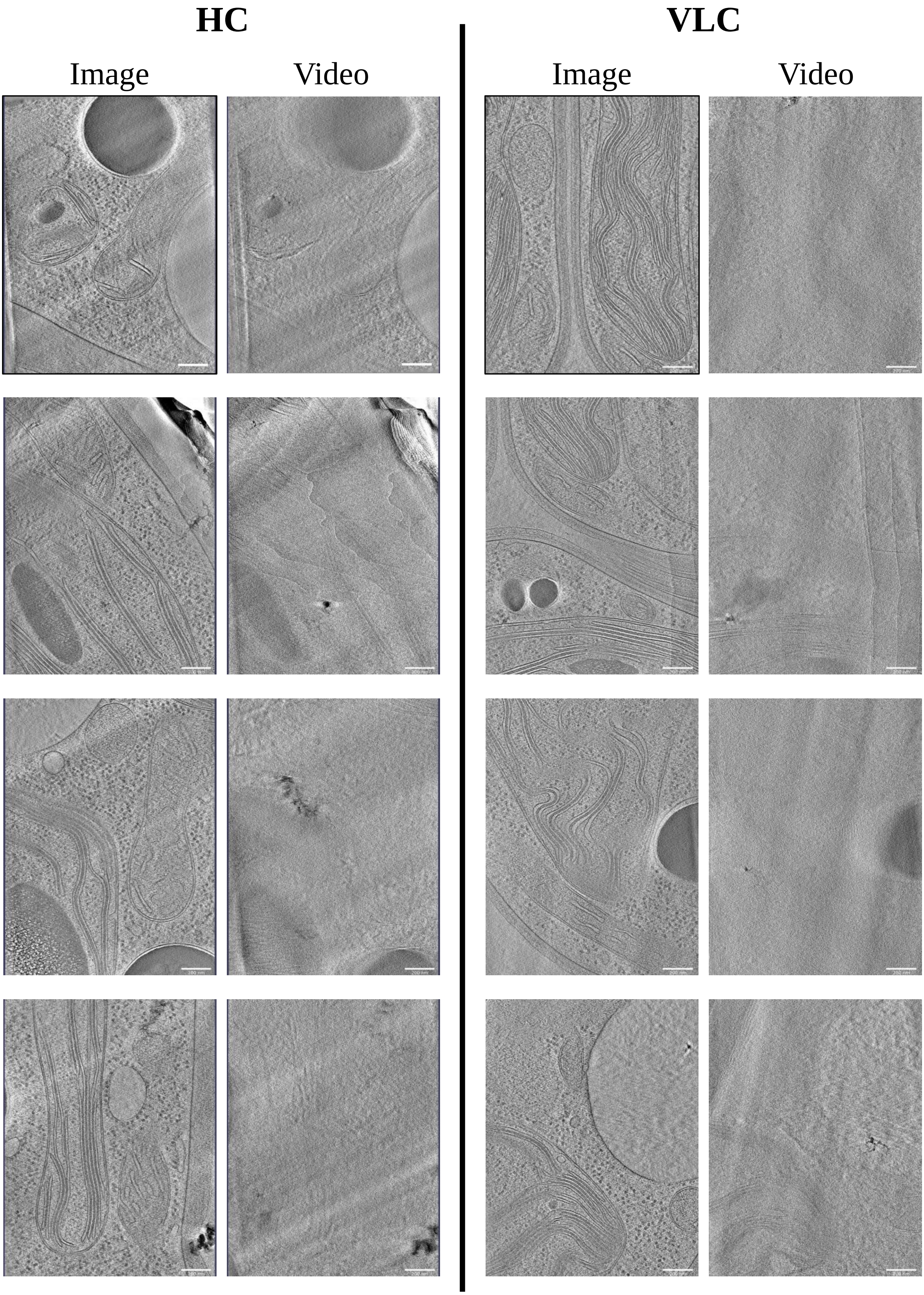

**Supplemental Figure 3: Kinetics of mitochondrial rearrangement during a shift from HC to VLC and VLC to HC. A.** Cells were immobilized under TP agar and acclimated to HC for 6 h before being exposed to either ambient air and HL (the initial LC attains VLC in HL). **B.** VLC grown cells were placed in the dark to allow the $CO_2$ level to rise (attaining HC). The upper rows of **A** and **B** are fluorescence signals from cross sections of the cell while the lower rows are images of the cortical region of the cells. The numbers in the left corner of the images (only shown in top row of each panel) are the times after the cells were shifted to the new conditions. Scale bar: 10 µm**.** Fluorescence microscopy images are representative of 2 experiments.

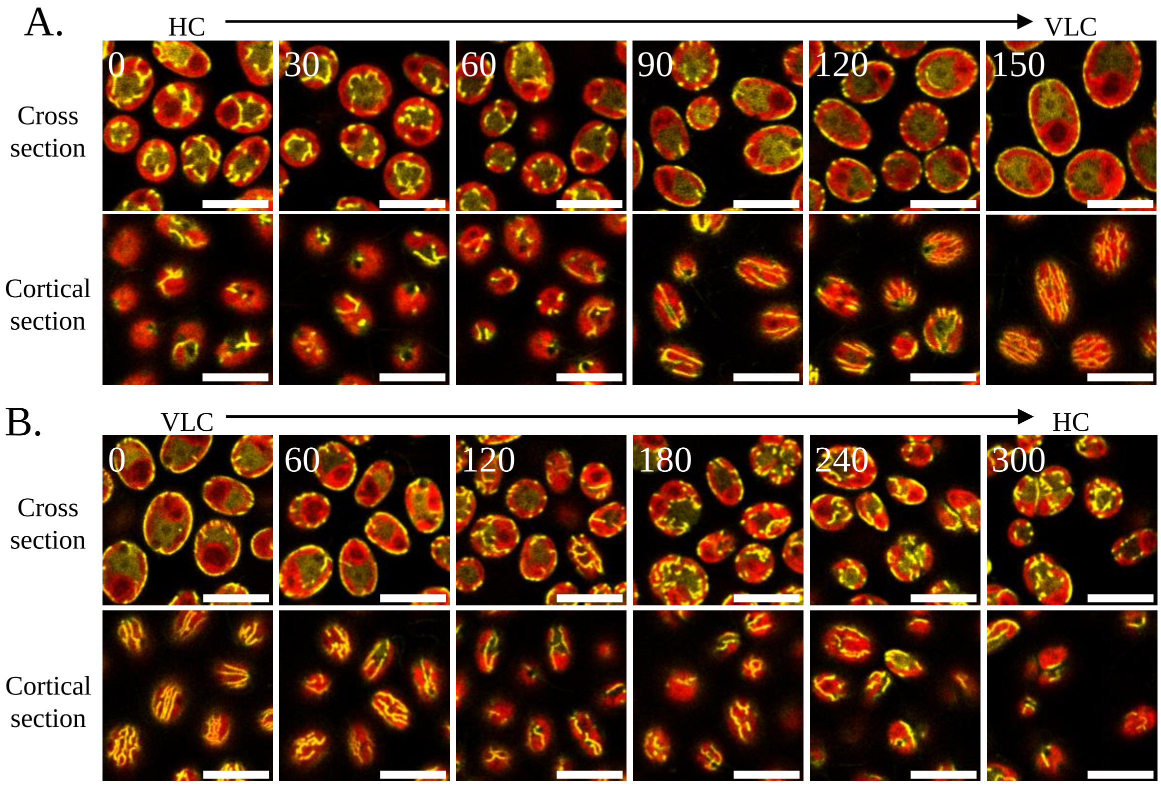

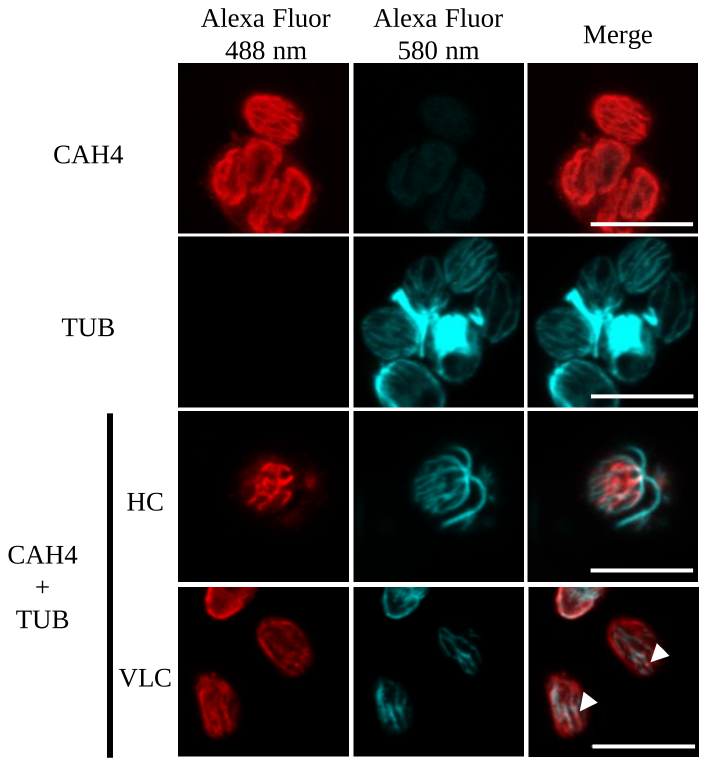


**Supplemental Figure 4. Immunofluorescence of mitochondria and microtubules in HC and VLC grown cells.** Cells were immuno-stained with antibodies directed against mitochondrial carbonic anhydrases (CAH4/5; red), α-Tubulin (TUB; blue) or both (CAH4 + TUB). Several slices were assembled into a projection of average intensity. Single antibody samples show absence of signal blead into the other channel. Immuno-staining with both antibodies to CAH4/5 and TUB was performed on cells incubated in HC and VLC for 24 h. Arrows show area of overlap between the mitochondria and microtubules. Scale bar: 10 µm. Fluorescence microscopy images are representative of 2 experiments.

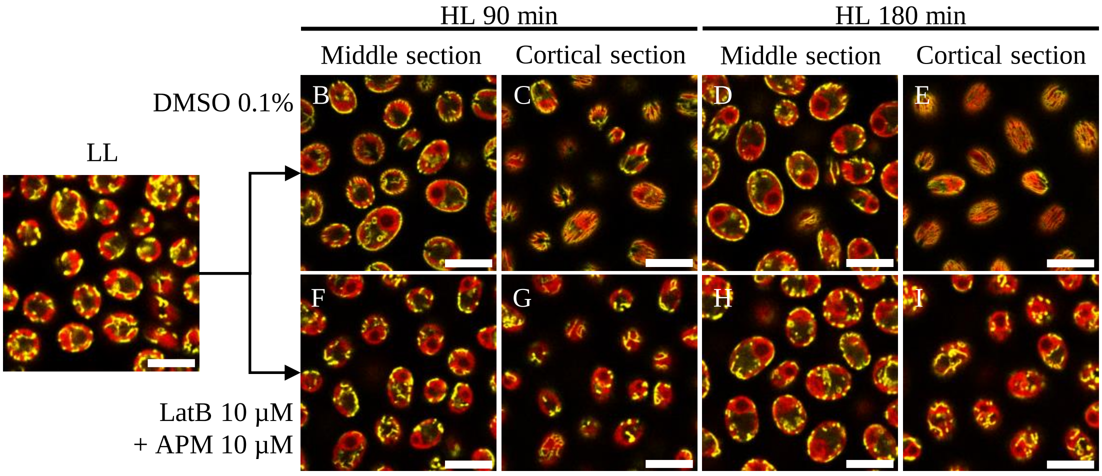


**Supplemental Figure 5. Effect of simultaneous LatB and APM treatment on mitochondrial dynamics.** Synchronized *nap1-1* cells were exposed to HL at the start of the day in the presence (LatB 10 µM + APM 10 µM) or absence of microtubule and actin inhibitors (control is 0.1% DMSO). Scale bar: 10 µm. Fluorescence microscopy images are representative of 2 experiments.

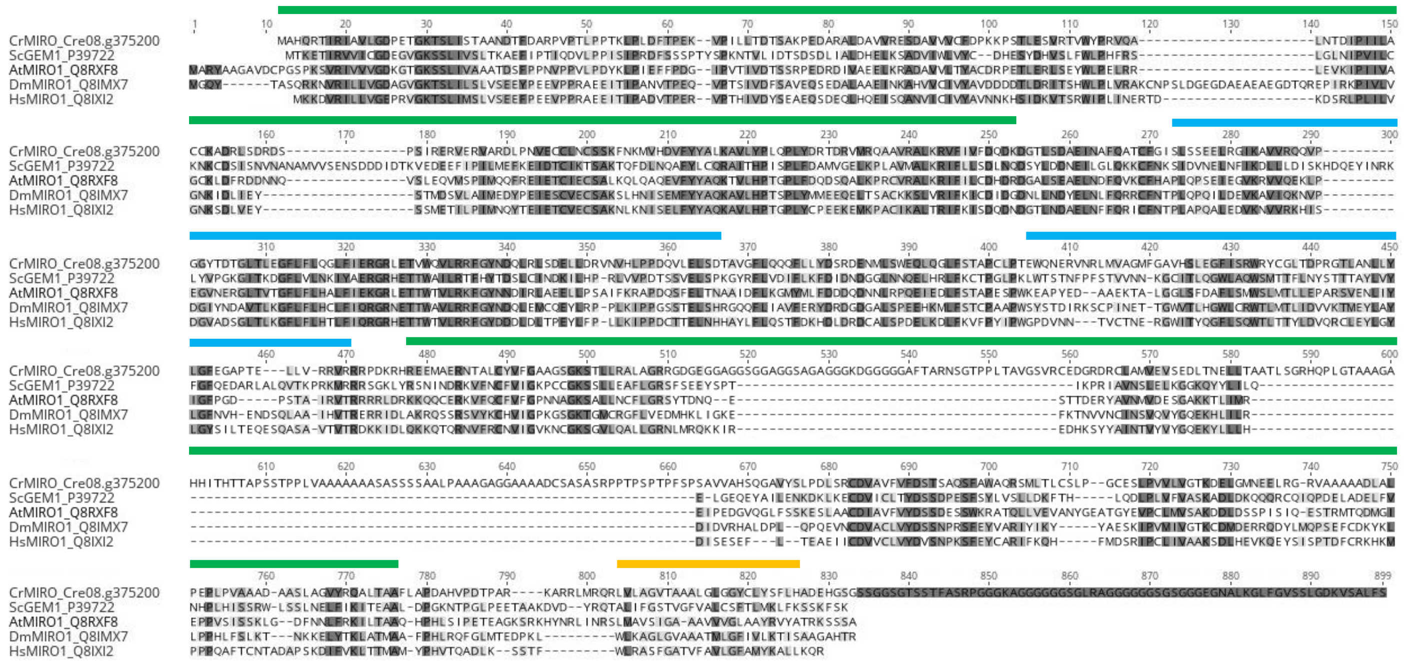


**Supplemental Figure 6: Alignment of MIRO proteins from *C. reinhardtii*, *S. cerevisiae*, *A. thaliana*, *D. melanogaster* and *H. sapiens*.** Alignment was performed using Clustal Omega. Access IDs are indicated next to the protein name. Chlamydomonas MIRO1 predicted domains are depicted above the sequence: GTPase domains 1 and 2 (green), EF hand domains 1 and 2 (blue) and C-terminal transmembrane domain (yellow). The current CrMIRO1 gene model includes large amino acid insertions in the predicted GTPase domain 2, and also has an extended C-terminal sequence.

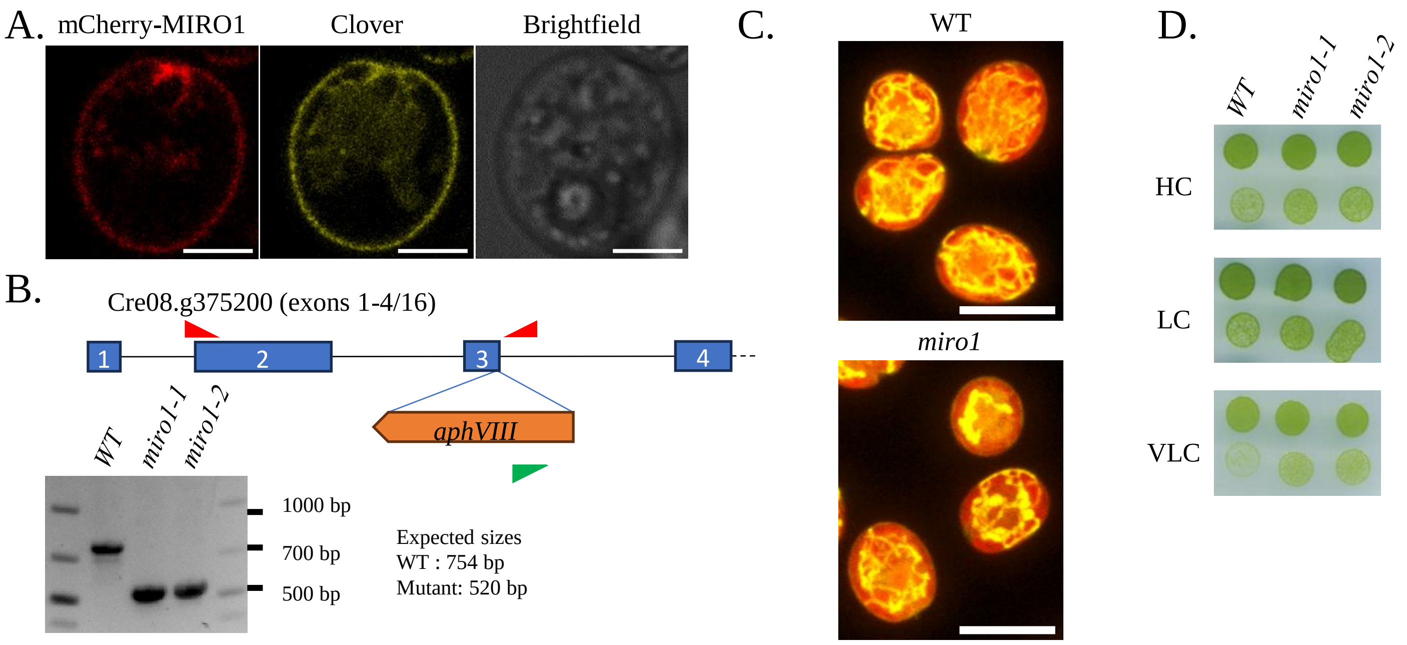


**Supplemental Figure 7**: **Phenotypic characterization of the *miro1* strains. A.** mCherry-MIRO1 and mito-Clover localization under VLC conditions. Scale bar: 2 µm. **B.** PCR showing the shorter product in the *miro1-1* and *miro1-2* strains. Because PCR conditions could not produce the fragment containing the full insertion cassette, we use a 3 primers approach with two primers annealing on both sides of the target CRISPR site in the third exon (red arrows) and one on the 3' side of the *aph7"* insertion cassette (green arrow). **C.** Mitochondria localization in HC grown cells of *miro1* and the parental WT strain. Scale bar: 10 µm. **D.** Growth test of the *miro1* strains and the parental strain CC-125. Cells were spotted on minimal medium and grown in chamber with $CO_2$-enriched atmosphere (HC), ambient air (LC) or $CO_2$-depleted air (VLC). Fluorescence microscopy images are representative of 2 experiments.

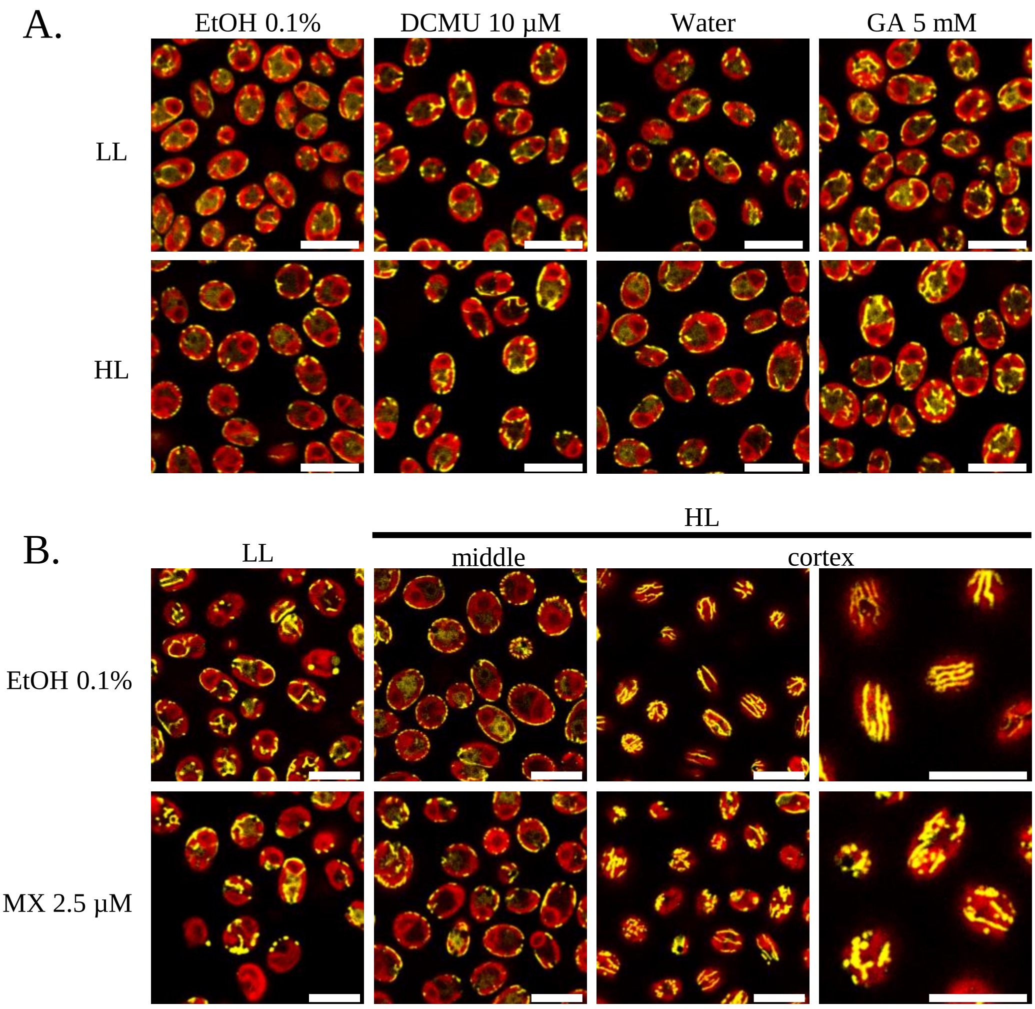


**Supplemental Figure 8**: **Effect of inhibitors on mitochondria relocation. A.** Effect of DCMU (10 µM) and Glycolaldehyde (GA; 5 mM): cells were mixotrophically grown in LL before testing peripheral relocation induced by HL exposure. Control samples (EtOH 0.1%; water) contained no inhibitor but were administered the same amounts of solvents added with inhibitor delivery (same final solvent concentration). Scale bars: 10 µm. **B.** Effect of 2.5 µM Myxothiazol (MX) on mitochondria relocation and cortical organization. Mixotrophically grown cells were exposed to HL in presence or absence of MX. Scale bar: 10 µm. Fluorescence microscopy images are representative of 2 experiments.

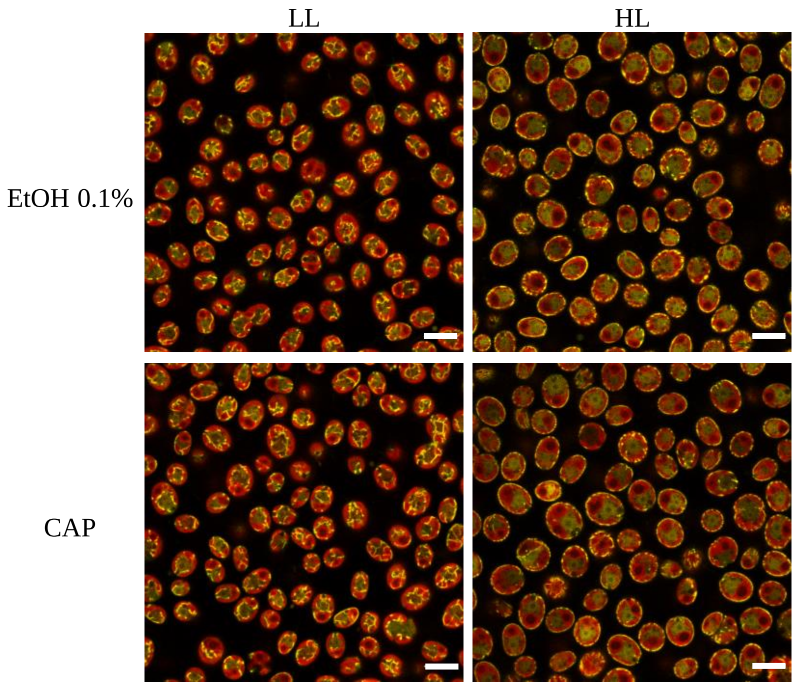


**Supplemental Figure 9**. **Effect of Chloramphenicol on mitochondria relocation**. Cells were grown in TAP LL and exposed to HL for 3 h in presence (250 µg $ml^{-1}$) or absence (EtOH 0.1%) of chloramphenicol (CAP). Scale bar: 10 µm. Fluorescence microscopy images are representative of 2 experiments.

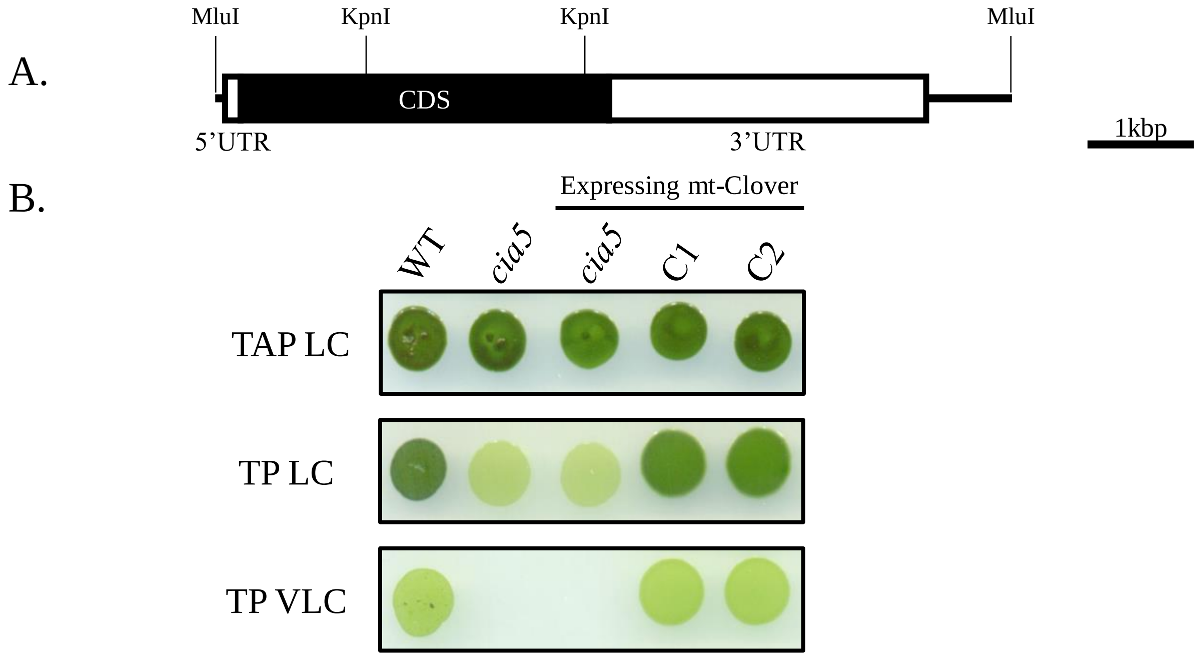


**Supplemental Figure 10**. **Complementation of the *cia5* mutant. A.** Map of the *CIA5* gene (Cre02.g096300) indicating restriction sites around the gene (MluI) used to extract it from the BAC vector PTQ9468, along with the positions of the KpnI sites within the gene, which were used as negative controls for complementation. **B.** Growth of the *cia5* mutants and complementing strains. Cell suspensions were deposited on agar medium (TAP or TP) and incubated for 5 days under LC or VLC conditions.

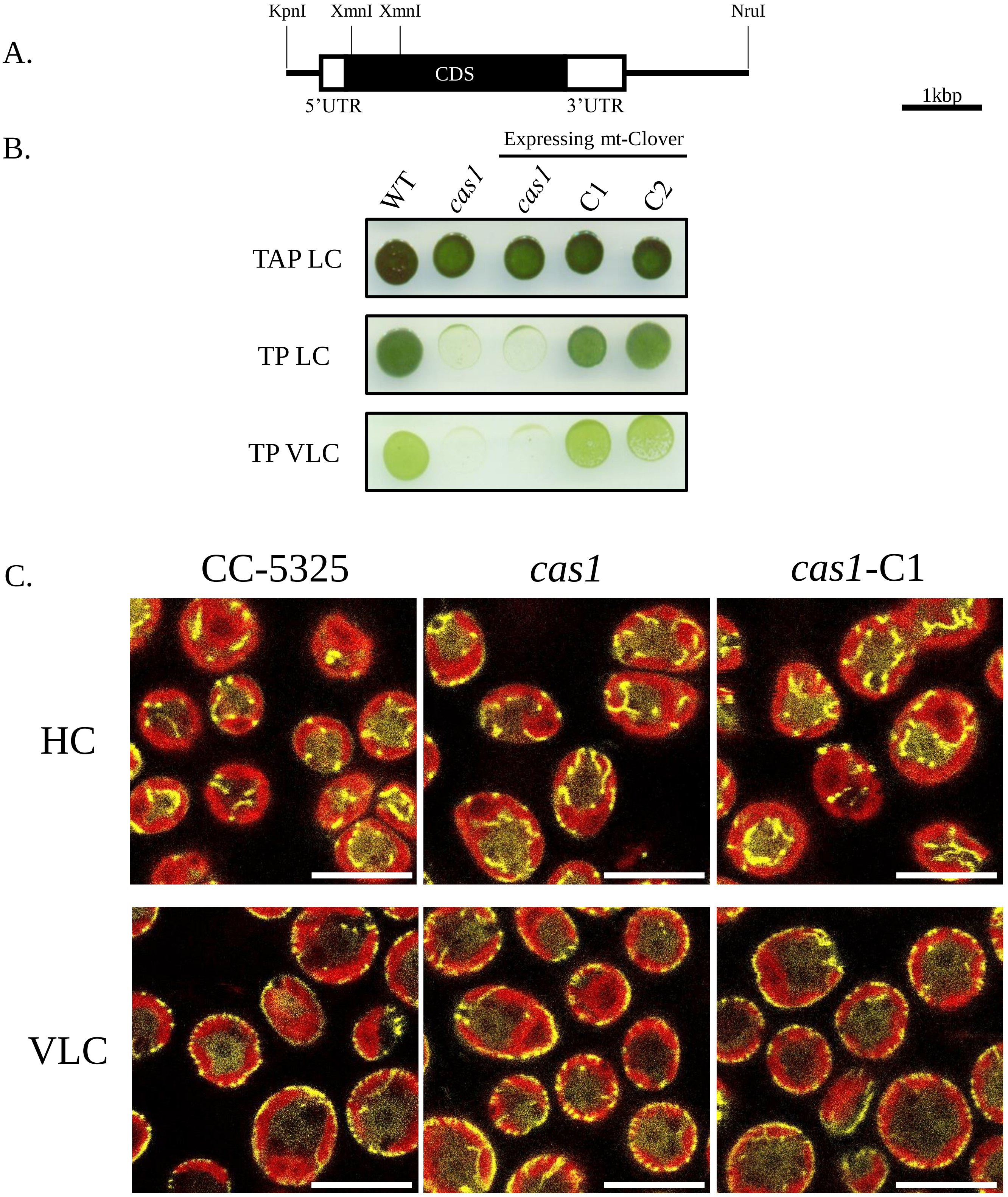


**Supplemental Figure 11**. Complementation of the *cas1* mutant. **A.** Map of the CAS1 gene (Cre12.g497300) indicating restriction sites around the gene (KpnI/NruI) used to extract it from the BAC vector PTQ9603 and those cutting inside (XmnI) used as negative control for the complementation. **B.** Growth test of the *cas1* strains and complementing strains. Cell suspensions were deposited on agar medium (TAP or TP) and incubated for 5 days under LC or VLC conditions. **C.** VLC dependent relocation of mitochondria in *cas1*. Wild-type (CC-5325), mutant (*cas1*) and complemented (*cas1*-C1) cells were grown photoautotrophically in HC and tested for mitochondrial relocation following 4 h of VLC treatment. Scale bar: 10 µm. Fluorescence microscopy images are representative of 2 experiments.

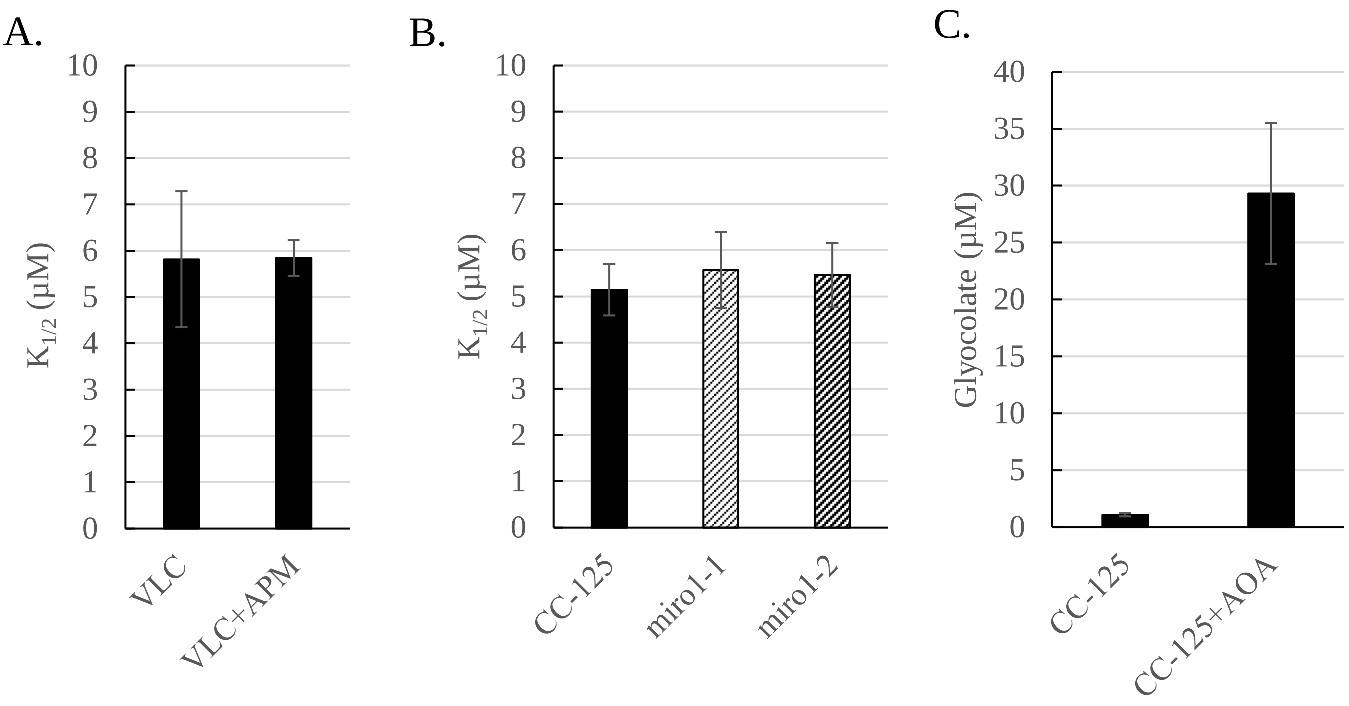


**Supplemental Figure 12**: **Affinity of *miro* mutants and inhibitor treated WT cells for inorganic carbon**. **A-B.** Ci affinity measurements of the effect of APM treatment (**A**) or absence of the MIRO1 protein (**B**) on the cell's affinity for Ci under VLC conditions (peripheral location of mitochondria). The data are the mean ± S.D (n=3). **C.** Glycolate content measured in the medium of VLC grown WT cells (CC-125) in the presence or absence of aminooxyacetate (AOA) for 24 h. The data are the mean ± S.D (n=3).

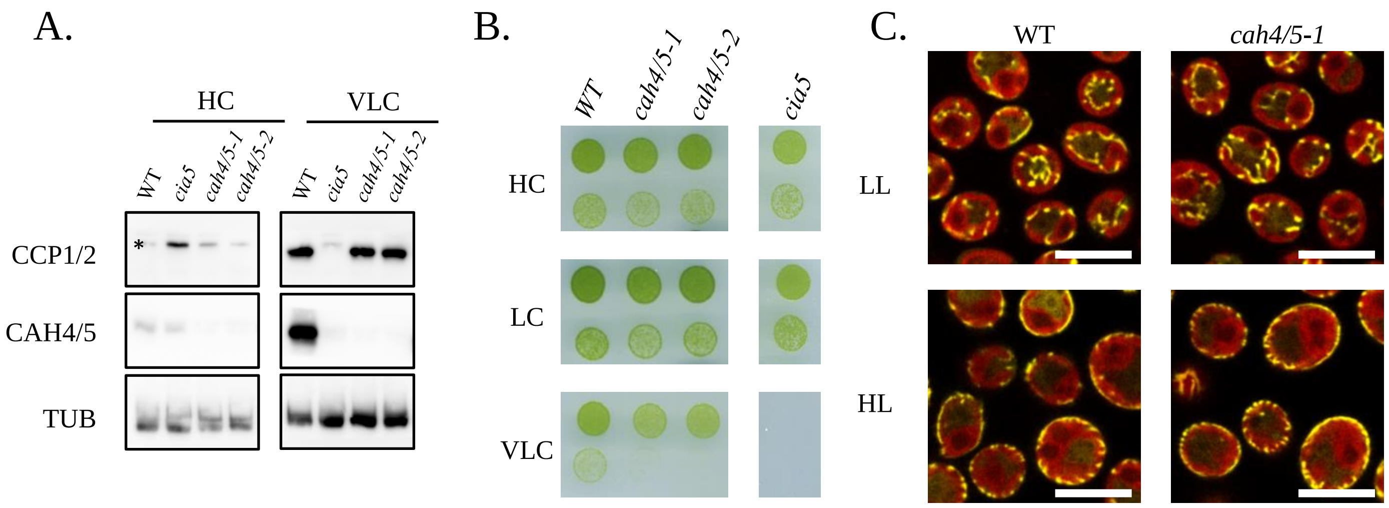


**Supplemental Figure 13**: **Characterization of the *cah4/5* mutant strains. A.** Immunodetection of CAH4/5 in HC and VLC grown cells. CCP1/2 antibodies were used as a positive control for CCM induction. *cia5* was used as a negative control of CCM induction. **B.** Growth test of the *cah4/5* strains and the parental strain CC-125. *cia5* is used as a control for absence of growth in VLC. Cells were spotted on minimal medium and grown in chamber with a $CO_2$-enriched atmosphere (HC), ambient/air levels of $CO_2$ (LC) or $CO_2$-depleted air (VLC). **C.** Mitochondrial relocation in the *cah4/5* double mutants. A mixotrophic culture of the *cah4/5* strain was exposed to HL for 3 h. The WT strain (CC-125) was used as a control. Scale bar: 10 µm. Immunoblot, growth test and fluorescence microscopy images are representative of 2 experiments.

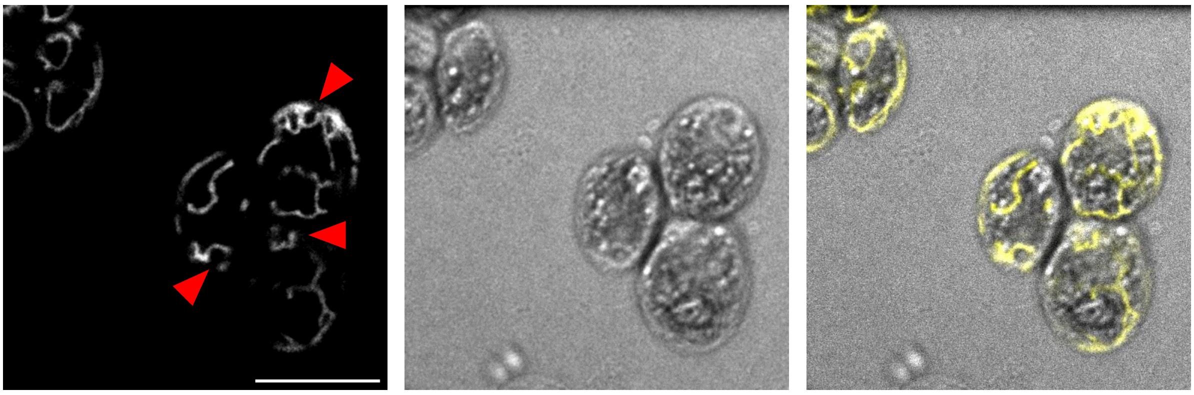

**Supplemental Figure 14: Timelapse of mitochondrial dynamics showing its association with the contractile vacuole.** Notice the rhythmic movement of mitochondria surrounding the contractile vacuoles near the apex of the cells (red arrowhead). The mitochondrial fluorescence is shown in white in the leftmost panel and in yellow in the rightmost panel. Scale bar: 10 µm.
